# Progenitor Diversity and Architecture of the Human Ganglionic Eminences Shaping the Basal Ganglia

**DOI:** 10.64898/2025.12.31.697063

**Authors:** Clara V. Siebert, Mengyi Song, Juan A. Moriano, Zhenmeiyu Li, Arantxa Cebrian Silla, Miranda Walker, Songcang Chen, Jennifer Baltazar, Lilian Gomes de Oliveira, Merut Shankar, Yuhan Xie, Pranav Suraparaju, Shaohui Wang, Qiuli Bi, Yajun Xie, Yuqi Ren, Miguel Turrero García, Li Wang, Guolong Zuo, Marten P. Smidt, Marco F.M. Hoekman, Corey Harwell, Jack Parent, John Rubenstein, Arturo Alvarez-Buylla, Arnold Kriegstein

## Abstract

The embryonic medial and lateral ganglionic eminences (MGE, LGE) are the principal sources of most neurons and glia for the basal ganglia. In primates, the MGE has a distinctive cytoarchitecture characterized by doublecortin enriched cellular nests (DENs), yet the architectonic organization underlying DEN formation, the molecular heterogeneity of ganglionic eminence progenitors and their lineage relationships, remain poorly understood. Here, using paired single-nucleus transcriptomics and chromatin accessibility profiling of the three GEs, we identify distinct progenitor populations, delineate their gene regulatory networks, and reconstruct their lineage trajectories. Live imaging reveals a unipolar outer radial glia-like population (GE-oRG) that undergoes mitotic somal translocation. Spatial transcriptomics identifies a distinct CRABP1+/ANGPT2+ domain within the MGE. Integrated spatial and electron microscopy demonstrates a periphery-to-center gradient of differentiation in the MGE. Leveraging DEN-forming MGE organoids derived from PCDH19 knockout human pluripotent stem cell lines, we identify the protocadherin, PCDH19, as a key regulator of DEN formation.

## Introduction

The basal ganglia, comprising the striatum, globus pallidus, subthalamic nucleus and substantia nigra, are subcortical regions essential for motor control^1^. During development, the lateral ganglionic eminence (LGE) serves as the developmental origin of the striatum acting and is the main source of striatal projection neurons, predominantly medium spiny neurons (MSNs), as well as olfactory bulb interneurons^2–5^. The medial ganglionic eminence (MGE) produces GABAergic interneurons destined for the striatum, globus pallidus and cortex, as well as projection neurons for the globus pallidus and other brain regions^6–12^. A third GE region, the caudal ganglionic eminence (CGE) produces mainly cortical interneurons and may contribute to other forebrain structures^6,7,13–17^. The three GE domains are distinguished by characteristic gene signatures: NKX2-1 marks the MGE, NR2F2 and SP8 characterize the LGE and CGE, and DLX family genes, that are core regulators of GABAergic development^14,18^, are differentially expressed across all GE subregions. The subthalamic nucleus and substantia nigra are not derived from the ganglionic eminences^19,20^ and are therefore not considered in this study. Overall, the ganglionic eminences serve as the principal source of GABAergic neurons that establish inhibitory circuitry throughout the telencephalon.

Recent single-cell transcriptomic studies have broadened our understanding of GE cell type diversity across species^21–29^, identifying regionally distinct progenitor populations and revealing conserved developmental programs. However, comparative analyses also suggest that primate GEs contain progenitor populations not found in rodents, including putative outer radial glia-like (oRG-like) cells and expanded intermediate progenitor (IPC) populations^21,25,30,31^, whose morphology, function, and molecular identity remain incompletely understood. Critical gaps also remain in understanding the full extent of GE progenitor heterogeneity, including underlying gene regulatory and epigenetic mechanisms.

The mammalian GEs exhibit a unique densely packed cellular organization that differs markedly from the layered architecture of the cortex^30–33^. This dense organization is accompanied by an expanded subventricular zone (SVZ) enriched with IPCs, features that distinguish primate GEs from their rodent counterparts^23,30,31^. In humans and other large-brain primates, but not mice, the MGE, but not LGE or CGE, exhibits a unique cytoarchitecture, with vimentin/nestin-expressing cells encircling DCX⁺ cell–enriched nests (DENs)^30,31^. This putative primate-specific MGE organization, may contribute to the expanded generation of interneurons in large-brained primates^30,31^. The differential expression of the protocadherin PCDH19 in neuroblasts and PCDH10 in surrounding nestin+ progenitor clusters has suggested a possible role for prorocadherins in DEN formation^30^. However, the spatial distribution of diverse progenitor populations within the MGE and their relationship to DEN formation remains poorly understood.

In this study, we investigated progenitor heterogeneity in the developing ganglionic eminences that produce and amplify many cell types that contribute to the basal ganglia and other brain regions. Using multiome sequencing, spatial transcriptomics and live imaging, we characterized molecularly distinct GE progenitor populations and their underlying gene regulatory networks. We identify a RG subtype, which we term GE-oRG, that shares hallmark morphological features and division behaviors with cortical oRG cells but exhibits distinct molecular features. Using spatial transcriptomics and electron microscopy (EM), we define the cellular organization of MGE DENs, uncovering a multi-level structure of RG fiber columns surrounding DENs that are further partitioned by IPCs and GE-oRGs. We propose this unique cellular cytoarchitecture as a driver of amplification for the neuronal output of the MGE. To investigate the mechanisms underlying DEN formation, we developed MGE-enriched GE organoids that form DEN-like structures, providing an *in vitro* platform to study human MGE cytoarchitecture. Using organoids derived from PCDH19-knockout and isogenic control hPSC lines, we demonstrate a critical role for PCDH19 in DENs formation. Lastly, by mapping the gene regulatory architecture of GE progenitors and their derived lineages, we link neuropsychiatric risk variants to specific cell types and developmental states. Our findings reveal that human GABAergic diversity emerges from a coordinated, multi-scale developmental strategy, where molecularly distinct progenitors occupy specialized niches that integrate lineage, spatial, and cytoarchitectural organization to amplify and orchestrate inhibitory neuronal production throughout the forebrain.

## Results

### Joint single-nucleus transcriptome and epigenome profiling of the developing human GE

To better define the cellular diversity and underlying transcriptional and epigenomic landscapes of human GE development, we collected LGE, MGE and CGE samples at gestational weeks (GW) 14 and 20 (GW14 and GW20). Leveraging the 10x Genomics single-nucleus multiome platform, we generated paired RNA-seq and ATAC-seq data, from which 20,319 high-quality nuclei were acquired for downstream analysis. At the single-nucleus level, we observed median values of 1,931 genes, 3,666 transcripts, and 8,980 ATAC peak fragments.

We integrated ATAC and RNA information using a weighted nearest-neighbor analysis followed by UMAP embedding and clustering. Drawing on established annotations^21,25,29,31,34,35^, we obtained 31 clusters with molecularly distinct cell types encompassing progenitors, neurons, and glia and clear segregation of LGE-, MGE-, and CGE-derived lineages (**Fig. 1A-D**). The progenitor compartment comprised clusters 0, 2, 5, 9, 11, 16, and 29, characterized by expression of one or more progenitor markers (*SOX2*, *VIM*, *ASCL1*, *DLX* family). A detailed analysis of this population is provided later. In the LGE-lineage, we identified multiple medium spiny neuron (MSN) lineages, including D1 MSNs (*PDYN*⁺ and *TSHZ1*⁺ lineages), D2 MSNs, and *FOXP1*/*ISL1*/*NPY1R*⁺ subpopulations. Within these LGE-derived lineages, some further split into multiple clusters, likely reflecting developmental or maturation states. The MGE contained diverse interneuron subtypes, including *SST*⁺ and *CRABP1*⁺ populations, while the CGE contributed *NR2F2*/*PROX1*⁺ interneurons. Subtype markers such as PV (*PVALB*), which emerges later in development^36^, and *VIP* were not yet robustly detected. We also identified cholinergic (*ChAT^+^)* neurons as well as ventromedial forebrain (VMF)**-**derived *GBX1*/*GABRA1*⁺, *ZIC1*/*ZIC2*⁺ neuronal populations. In addition to neuronal lineages, glial classes were also present, including astrocytes, oligodendrocyte precursor cells (OPCs), and microglia.

**Figure 1.**
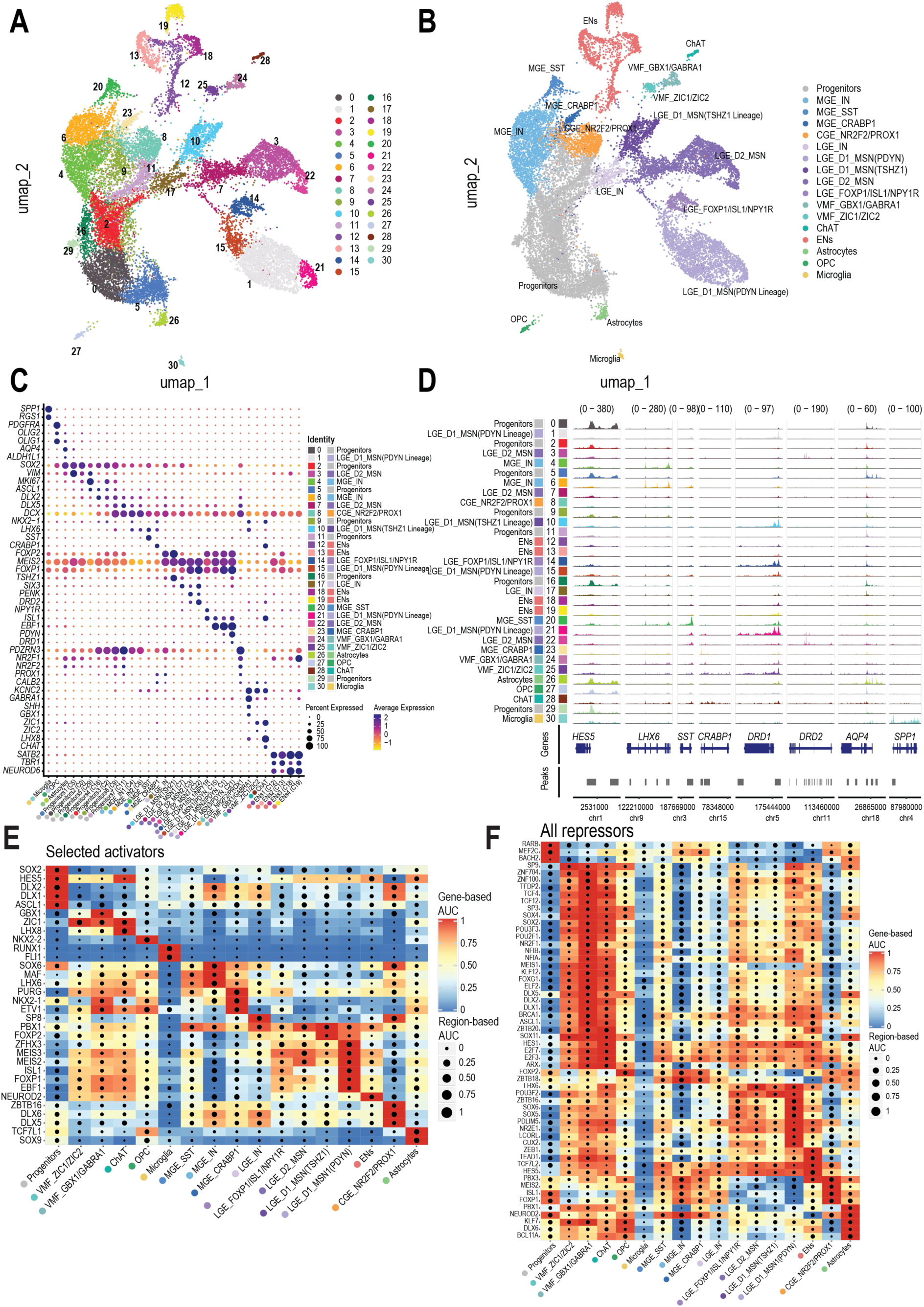
Multiome sequencing reveals cell type diversity in the developing ganglionic eminences. **A**-**B**) UMAP visualization of merged single-nucleus datasets from GW14 and GW20 LGE, MGE, and CGE samples, colored by cluster IDs (**A**) or annotations (**B**). **C**) Dot plot showing the mean expression levels (color) of marker genes and the percentages of cells in which marker genes are expressed (size) in each cluster. **D**) Aggregated chromatin accessibility profiles at locus of signature genes across cell type. **E, F**) Heatmaps showing the region-based AUC scores and gene-based AUC scores of selective activators (**E**) and all repressors (**F**) eRegulons across cell types. Cell types are ordered according to their gene-expression similarity.

Distinct patterns of open chromatin shape both cellular identity and developmental potential^37–41^. To understand the regulatory landscape defining diverse cell lineages in the developing human GEs, we examined differentially accessible regions (DARs). DARs revealed distinct regulatory programs across progenitor states and lineage progression (**Fig. 1D, Supplemental Fig. 1, 2**). Progenitor-wide DARs (clusters 0, 2, 5, 9, 11, 16, 29) highlighted biological processes related to stem cell maintenance, proliferation, glial fate, and Notch signaling (**Supplemental Fig. 2A, F,** Data S1). As the progenitor lineages progressed (**Fig. 1A, Supplemental Fig. 2A**), they acquired DARs associated with GABAergic differentiation and olfactory bulb development (**Supplemental Fig. 2F, Data S1**), indicating emerging lineage competence. Lineage-specific DARs further delineated GABAergic regulatory programs: MGE-derived lineage (clusters 4, 6, 20, 23 in **Fig. 1A**) exhibited accessibility near genes governing GABAergic interneuron differentiation, migration, and maturation (**Supplemental Fig. 2B, D, Data S1**). For example, DARs in the *SST^+^* interneurons were highly enriched for proximity to genes related to characteristic of cortical interneuron programs. LGE-derived lineages, in turn, showed enrichment for chromatin regions associated with olfactory bulb interneuron development, habituation, and dopaminergic pathways, implicating their roles in sensory and reward-related processes (**Supplemental Fig. 2C, E, Data S1**).

Transcription factors (TFs) orchestrate lineage progression in the GEs by regulating gene expression programs in progenitors and differentiating neurons. Because motif enrichment alone does not indicate functional TF activity, we used SCENIC+^42^ to integrate single-nucleus ATAC and RNA data with motif discovery to infer enhancer-driven regulons (eRegulons) in the development of inhibitory neurons. Using this approach, we identified 555 eRegulons, with 389 transcriptional activators and 166 repressors, targeting 45,013 regions and 6,048 genes (**Fig. 1E, F**). Activator gene regulatory network (GRNs) analysis recovered known master regulators of GE progenitors (*SOX2*, *HES5*, *ASCL1*, and DLX family genes) and regional TFs (**Fig. 1E**): MGE: *NKX2-1*, *LHX6*, LGE: *ISL1*, *EBF1*, *MEIS2*, *SP8* and CGE: *ZBTB16, SP8*. Notably, the MGE-CRABP1+ lineage showed the strongest enrichment of the NKX2-1–GRN among MGE-derived populations, suggesting a potential striatal-biased fate, as *NKX2-1* expression is maintained in striatal interneurons but downregulated in cortical interneurons^43^. Our analysis also uncovered activator programs that are less well characterized in this context, including PURG in the MGE-CRABP1+ lineage and ZFHX3 in the LGE MSN lineage (notably the D1 MSN PDYN subtype). In contrast to activators, which form highly cell-type specific GRNs, repressor-mediated GRNs acted more broadly across diverse cell types (**Fig. 1F**). This widespread activity likely reflects their essential role in safeguarding cell identity by continuously suppressing alternative fate programs, thereby preserving lineage fidelity during neural development.

### GE-derived cell types are associated with neuropsychiatric diseases

Given that most neuropsychiatric genome-wide association study (GWAS) risk variants fall within non-coding regulatory elements^44–46^, we mapped GWAS variants associated with neuropsychiatric disorders onto our single-nucleus chromatin accessibility profiles. We examined a variety of neuropsychiatric disorders, including attention-deficit/hyperactivity disorder (ADHD), autism spectrum disorder (ASD), bipolar disorder (BD), major depressive disorder (MDD), obsessive-compulsive disorder (OCD), and schizophrenia (SCZ)^47–59^. We employed SCAVENGE to systematically link GWAS variants to their cellular contexts (**Supplemental Fig. 3**). LGE and MGE lineages showed enrichment across multiple neuropsychiatric traits, highlighting lineage-specific vulnerability. Some key highlights emerged: LGE-derived MSNs are associated with ADHD (D1 TSHZ1⁺, **Supplemental Fig. 3A, G**), pointing to a striatal developmental association and potential links to dopaminergic pathways implicated in ADHD^60–63^ and MDD^64^ (broad MSN enrichment, **Supplemental Fig. 3D, G**). MGE-derived SST⁺ interneurons are strongly linked to SCZ (**Supplemental Fig. 3 F, G**) aligning with their established roles in cortical inhibition central to schizophrenia pathogenesis^65–70^. Additionally, a VMF-derived GBX1⁺/GABRA1⁺ population, likely globus pallidus–destined^29,71^, also associates with MDD (**Supplemental Fig. 3D, G**), consistent with accumulating evidence in the literature^72,73^. Collectively, these results support that some genetic risk for certain neuropsychiatric diseases is cell-type specific within the ventral forebrain.

### Molecular heterogeneity of human Radial Glia in the GEs reflects regional differences

To investigate how the diversity of GE-derived lineages emerges, we focused on their progenitors. We subclustered the defined progenitor and glia compartments (clusters 0, 2, 5, 9, 11, 16, 26, 27, 29 in **Fig. 1A**). We identified six major clusters: RG subtypes or states (*HES5⁺/VIM⁺/SOX2⁺/PDGFRB⁺*), IPCs (*ASCL1⁺/DLX1⁺/DLX2****⁺***), neuroblast populations (*DCX⁺/DLX5****⁺/****DLX6**^+^***), where neuroblasts refer to IPC daughter cells strictly committed to a single neuron fate, along with cycling clusters (*MKI67⁺/TOP2A****⁺***), oligodendrocyte precursor (OPC; *OLIG2*) and astrocyte progenitor (APC; *ALDH1L1*) clusters (**Fig. 2A, B, Supplemental Fig. 4A, B, 5**). To minimize biases from dissection variability or migrating cells, we grouped transcriptionally similar clusters into broader classes and annotated them using known markers^21,25,29,31,34,74^ (**Fig. 2A, Supplemental Fig. 5, 6A**). RG and IPCs assignments largely agreed with dissection labels (**Supplemental Fig. 4C, D**). This resulted in the following groups: L/CGE RG, MGE RG, L/CGE IPC (early/late), MGE IPC (early/late), MGE neuroblast, L/CGE neuroblast, LGE neuroblast, and CGE neuroblast (**Fig. 2A, Supplemental Fig. 6A**).

**Figure 2.**
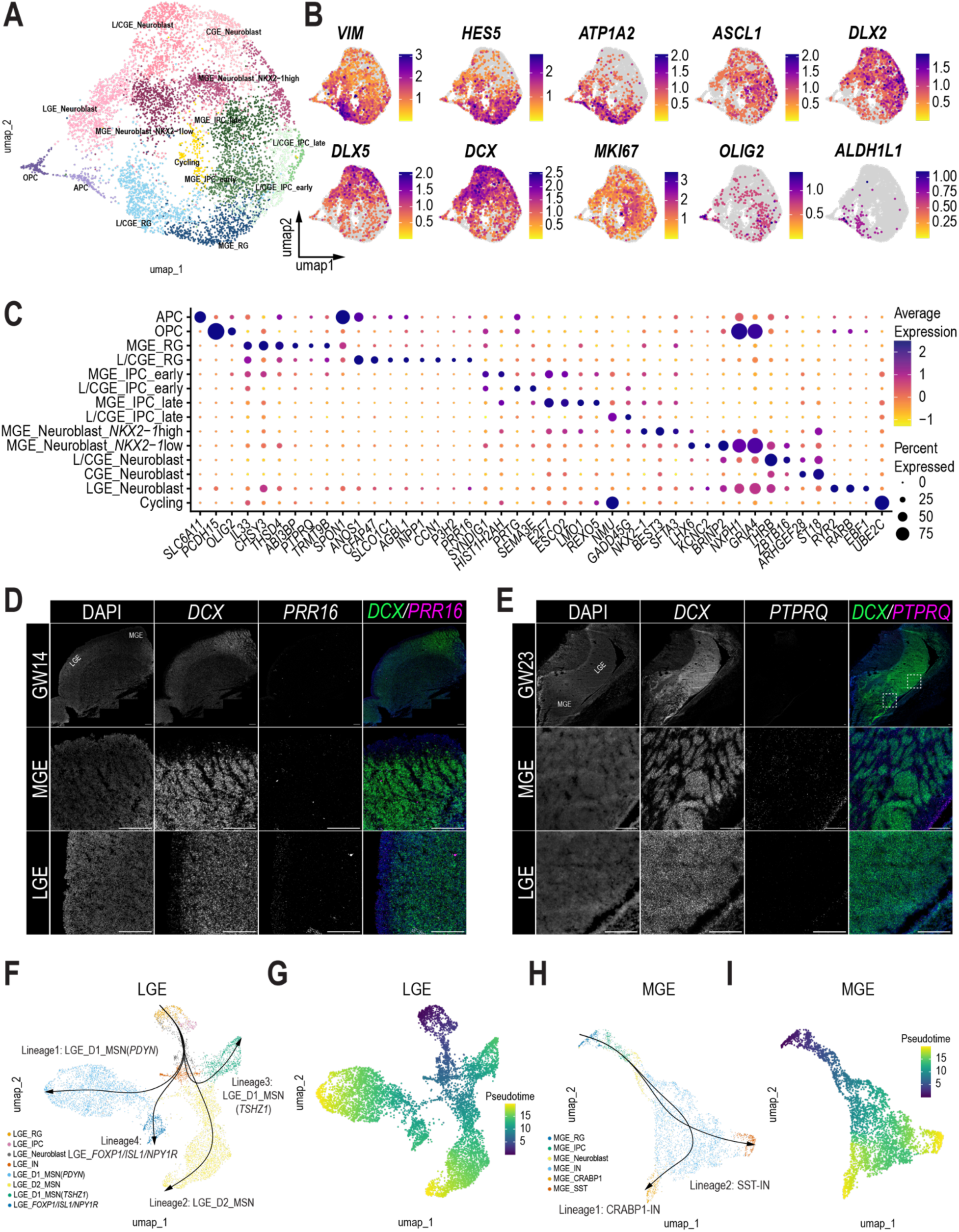
Progenitor diversity and lineage in the developing GEs. (**A, B**) UMAP visualization of progenitors from merged single-cell datasets from GW14 and GW20 LGE, MGE, and CGE samples, colored by cluster IDs (**A**) or expression level of marker genes (**B**). **C**) Dot plot showing the mean expression levels (color) of marker genes and the percentages of cells in which marker genes are expressed (size) in each cluster. **D, E**) RNAscope of LGE/CGE-RG gene *PRR16* in human GW14 sample (**D**) and MGE-RG gene *PTPRQ* in human GW23 sample (**E**). Scale bars, 100 μm. **F**, **G**) UMAP plots of cells from LGE neuronal lineages, colored by cell types (**F**) or pseudotime (**G**). **H**, **I**) UMAP plots of cells from MGE IN lineages, colored by trajectory clusters (**H**) or pseudotime (**I**).

RG displayed transcriptionally distinct, regional biased subtypes across the LGE, MGE, and CGE (**Fig. 2C, Supplemental Fig. 5B**). All RG expressed canonical markers (*VIM, SOX2, PDGFRB*), reflecting a shared progenitor identity. Pan-RG markers (*ATP1A2, PTPRZ1, TNC*) showed broad expression across progenitor zones in the MGE and LGE, with RG in the MGE notably enriched around DENs (**Supplemental Fig. 5, 6A**). Regional specification was marked by *NKX2-1* and *SOX6* in MGE RG (**Supplemental Fig. 5B**), *NR2F2* in LGE/CGE RG, and *PAX6* in the dorsal LGE and CGE (**Supplemental Fig. 5B**), reflecting early molecular diversification among RG populations. Consistent with previous studies^21^, additional regionally enriched genes—including *RIC3* and *MBIP* in MGE RG and *NTRK2* and *GLI3* in L/CGE RG—further contributed to the molecular separation across regional domains (**Supplemental Fig. 5B**). We also identified novel regional specific markers (**Fig. 2C**), including *PTPRQ* in MGE RG (**Fig. 2C, D**) and *PRR16* in LGE RG (**Fig. 2C, E**), which were validated by RNAscope. Overall, this analysis provides a high-resolution view of RG regionalization, revealing previously unrecognized molecular distinctions that refine our understanding of GE RG diversity.

### IPC and neuroblast diversity in the human GEs

IPCs have been proposed as the main transit-amplifying progenitors in the human GE^21^, expanding the neuronal output from this region. In our progenitor-enriched multiome dataset, IPCs formed an early-to-late differentiation axis across regions: early IPCs expressed relatively higher *SOX2*, *EGFR*, *HES5*, and *VIM*, consistent with residual RG-like features, whereas late IPCs displayed reduced levels of these genes and increased levels of *DLX1*, *DLX2*, *ASCL1*, *DLL1*, and *DLL3*. Consistent with the known sequential activation of DLX genes^75,76^, we observed a progressive shift in DLX family expression, with *DLX1/2* defining late IPCs and early NBs and *DLX5/6* marking the transition to *DCX⁺* neuroblasts (**Fig. 2B, Supplemental Fig. 5A, 6A**). While these features were broadly conserved across the GE, molecular differences emerged between regions (**Fig. 2C, Supplemental Fig. 5, 6A**). Cell-cycle analysis showed that RGs exhibited substantial G1-phase enrichment, while IPCs were predominantly in S or G2/M phases (**Supplemental Fig. 4E, F, 6B**), consistent with their role as transit-amplifying progenitors. This pattern aligns with strong Ki67 and ASCL1+ IPC enrichment in the iSVZ (**Supplemental Fig. 12D, E**). Interestingly, a subset of MGE early IPCs exhibited *OLIG2 and high EGFR* expression, resembling cortical tri-potent IPCs that can produce OPCs, interneurons, and astrocytes^77^. Immunostaining confirmed the presence of EGFR+/OLIG2+ progenitors, which were concentrated in the iSVZ and oSVZ (**Supplemental Fig. 6E**) indicating a similar Tri-IPC-like population may exist within the GEs but their lineage remains unclear.

Neuroblast populations across the human GE (LGE, MGE, and CGE) were defined by high DCX expression with DLX5/6, spanning late-stage progenitors to post-mitotic newborn neurons. LGE and CGE neuroblasts begun to express *SCGN* and *SP8,* along with some novel markers expressed in subtypes, such as *THRB* and *ARHGEF28*. MGE neuroblasts segregated into *NKX2-1*-high (Neuroblast1–3, *BEST3*^+^/*SFTA3^+^*), and *NKX2-1*-low populations (*LHX6^+^* Neuroblast8–9, **Fig. 2C; Supplemental Fig. 5B**). The *NKX2-1*-low populations were also enriched for genes identified in previous studies such as *NXPH1* and *PDZRN4*^21^ and featured novel markers such as *KCNC2* and *BRINP2*. Some neuroblasts were actively engaged in G2/M or S phase (**Supplemental Fig. 4F, 6B**), most pronounced in MGE neuroblasts which are likely the highly proliferative neuroblasts previously identified in DENs^30^. While the precise relationship between these MGE neuroblasts is not yet clear, they likely represent either sequential maturation stages or early divergence toward striatal versus cortical fates.

Having defined the diversity of GE progenitors, we next sought to connect them with basal ganglia lineages. Trajectory analysis using eRegulon revealed four divergent LGE trajectories: D1_MSN (PDYN), D1_MSN (TSHZ1), D2_MSN, and LGE_FOXP1/ISL1/NPY1R (**Fig. 2F, J**, **Supplemental Fig. 7A, F**), corresponding to basal ganglia projection neurons. At the same time, MGE progenitors diverged into two major lineages (*SST⁺* and *CRABP1⁺*; **Fig. 2H, I**, **Supplemental Fig. 8B, G**). NKX2-1 dynamics segregated MGE fates^43^: *CRABP1^+^* cells retained *NKX2-1*, suggesting a striatal fate, while *SST^+^* cells downregulated *NKX2-1*, indicating a cortical fate (**Supplemental Fig. 7C**). Notably, this bifurcation also displayed a temporal difference: *CRABP1*⁺ interneurons were predominantly represented at GW14 samples, whereas *SST⁺* cells were more abundant at GW20, suggesting that striatal interneurons are produced earlier than cortical interneurons (**Supplemental Fig. 7D**). The MGE also generates PV⁺ interneurons^14^. Rather than identifying a distinct PV trajectory, we identified gene signatures characteristic of PV neuroblasts and immature neurons^35^ enriched within the *CRABP1*⁺ lineage (**Supplemental Fig. 7E**) consistent with the presence of CRABP1⁺/PV⁺ cells in the adult human cortex^21^. Temporal dynamics of eRegulon activity defined ten modules in the MGE and ten in the LGE, each following distinct temporal patterns (**Supplemental Fig. 7A, B**). For example, in the MGE, modules transitioned from early BMP signaling to neuronal/glial specification and GABAergic interneuron differentiation (**Supplemental Fig. 7F, G**) highlighting the structured regulatory logic guiding GE lineage progression.

Together, these results demonstrate that diverse GE progenitors give rise to distinct GABAergic inhibitory neuron lineages through sequential amplification and differentiation.

### Mitotic GE-oRG cells constitute a large proportion of the GE progenitor pool

The presence of oRG-like cells in the developing GE has been suggested by previous studies but defined characterization of oRG-like cell biology and division behaviors across GEs is still lacking. We first assessed expression of the canonical oRG marker *HOPX*^78^ in our multiome dataset and assessed protein expression by immunostaining. Interestingly, *HOPX* was only expressed in a subpopulation of L/CGE RG, but not in MGE RG (RG1, **Supplemental Fig. 5B; Supplemental Fig. 10A, B)**, suggesting *HOPX* may be a regional oRG identity marker rather than a universal oRG marker.

To functionally characterize division behaviors in the developing GEs, we performed live imaging of vibratome slices of human MGE, LGE and CGE (GW13–14 and GW17–19) labeled with Ad-CMV-GFP. Adjacent sections were immunostained for NKX2.1, SP8, and NR2F2 to confirm regional identity (**Fig. 3A; Supplemental Fig. 10C**). Across all GEs, dividing non-ventricular RG-like cells with a prominent basal process underwent MST consistent with oRG identity (**Fig. 3B-D**). We termed these cells as ‘GE-oRG’ (**Fig. 3B**) reflecting their morphological and behavioral similarity to cortical oRG^31^. ASCL1^+^ immunostaining (**Fig. 3E, F**) demonstrating that these cells were not IPCs (**Fig. 3E, F).** Quantification of division types revealed temporal and regional differences. In the MGE at GW13–14, MST divisions predominated compared to IPC divisions (63.5% vs 25.0%, p = 0.0419) (**Fig. 3C, D**), consistent with a predominance of founder cell expansion at this relatively early stage of development. In the CGE, IPC-like (32.5%, p = 0.0491) and oRG-like (58.5%, p = 0.046) divisions were significantly more frequent than RG-like interkinetic nuclear migration (INM events, 9.0%). By GW17–19, cells undergoing IPC-like mitotic rounding prior to division increased in the MGE (63.0%, p = 0.0305) and LGE (45.0%, p = 0.0123), marking a shift toward differentiation, whereas proportions remained stable in the CGE (44.3%, **Fig. 3C, D**). MST translocation distances were longest in the CGE (65.2 µm) compared with the MGE (46.3 µm) and LGE (47.8 µm; p < 0.025) (**Fig. 3G; Supplemental Fig. 10D-F**), and interestingly cortical MST distances (average of 57µm)^79^ lay in between MGE/LGE and CGE. MST directionality was largely random (**Supplemental Fig. 10J**), consistent with the non-radial orientation of GE fibers^30,31^. Most MST divisions were orthogonal (60–90°) to the main process (71–95% across regions and ages) (**Supplemental Fig. 10G-I**), resembling cortical oRG behavior ^80,81^.

**Figure 3.**
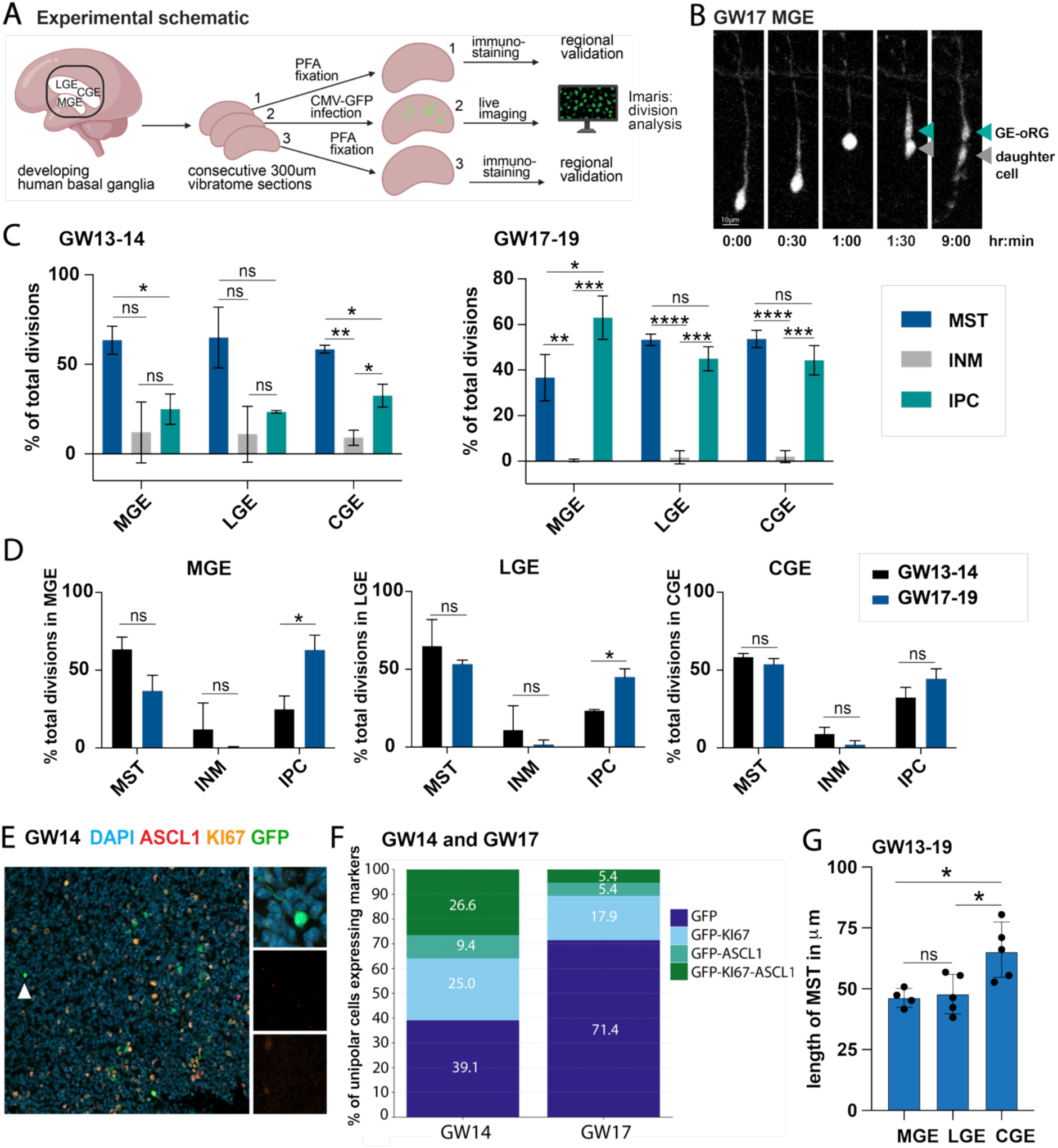
Live Imaging of human primary developing MGE, LGE and CGE. **A)** Experimental schematic of organotypic slice culture preparation and regional validation (created using BioRender.com). **B)** Example of MST in GW17 MGE sample with GE-oRG cells (green) and daughter cell (grey). Scale is 10μm. **C, D)** 815 divisions were analyzed. Quantifications of division modes in GEs across GW13-14 (n=2) and GW17-19 (n=3). **E)** Immunostaining of live imaged sections of GW14 MGE. **F)** Quantifications of cell type marker expression (immunostaining) in cells with one dominant process in GW14 (n=1) and GW17 (n=1) live imaged sections. **G)** MST length at GW13-19 (n=4-5, 415 MST events analyzed) in μm.

Together, these findings identified oRG-like progenitors within all three human GEs and revealed region- and stage-specific differences in division mode, translocation length, and MST division plane, suggesting distinct regulatory mechanisms of progenitor dynamics within the ventral telencephalon.

### Conserved and divergent RG programs between pallial and subpallial forebrain regions

To further assess shared and divergent molecular features of human RG, including oRGs, across the cortical (dorsal) and GE (ventral) regions, we compared their transcriptomic and epigenomic landscapes using publicly available cortical datasets^77^ (**Supplemental Fig. 8, 9, DataS2**). Transcriptomically, cortical and GE RG shared core markers (*VIM*, *SOX2*, *PTPRZ1*, **Supplemental Fig. 8E-J**), but showed clear regional divergence: GE RG expressed *NKX2*-*1*, *NR2F2-AS1*, *SIX3*, while cortical RG expressed *TFAP2C* (**Supplemental Fig. 8E-N**). We also identified novel region-specific genes: *TMEM132B*, *PLPPR1*, *CNTN3*, and *CDON* which were strongly enriched in cortical RG, while *PLXDC2*, *HES5*, *BRINP3*, *GHR*, and *GREB1L* showed significantly higher expression in GE RG. Further comparisons of GE RG subtypes (RG1-5) to the three established cortical RG subtypes (ventricular RG [vRG], outer RG [oRG], and truncated RG [tRG]^32,79,82^) (**Supplemental Fig. 8A-D, F-J**) showed that MGE RG4 and L/CGE RG1 were closest to cortical oRG. oRG markers common across cortex and GEs included *VIM* and *TNC*, while *FGFR2* and *LDB2* were enriched in the MGE (RG4), and *NTRK2* in the L/CGE (RG1). At the chromatin level (**Supplemental Fig. 9**), most accessible peaks were shared across regions (**Supplemental Fig. 9F-J**, e.g. *TTYH1*^21,83–85^). However, divergent accessibility patterns appeared to arise from two main sources. First, regional differences were evident with cortical RG showing accessibility near *PAX6* and *FEZF2*, and GE RG near *OLIG2*, *GSX2*, and *NKX2-1*. Second, epigenetic priming pre-configured cortical RG for excitatory neurogenesis (e.g., NEUROD2) and GE RG for GABAergic and olfactory bulb differentiation, preceding transcriptional divergence (**Supplemental Fig. 9E-L,** binomial test, Data SX). Comparison of cortical and GE RG subtypes showed limited overlap of accessible peaks despite conserved lineage relationships mirroring transcriptome similarities (**Supplemental Fig. 8D, 9A–D, F–J)**. The limited overlap of marker peaks between ventral and dorsal regions likely reflects epigenomic priming^40,86^, which encodes progenitor competence that magnifies differences beyond those detectable at the transcriptomic level (**Supplemental Fig. 8L, 9L-N**).

Together, these analyses suggest that ventral and dorsal progenitors rely on shared core regulatory machinery but diverge through lineage-specialized gene networks that drive distinct developmental outcomes.

### Spatial transcriptomics of the MGE and LGE provide insights into cellular architecture

Beyond molecular identity, neural progenitors and differentiating neurons function within three-dimensional tissue microenvironments, where spatial organization and cell-cell interactions guide developmental outcomes. The human MGE exhibits a putative primate-specific cytoarchitecture with mutually exclusive domains of DENs surrounded by VIM⁺ progenitor clusters^30,31^. To better understand the spatial and cellular organization of this distinctive structure, we performed spatial transcriptomic profiling of the human MGE, LGE and caudate using a custom panel of 433 GE-enriched genes and multiplexed error-robust fluorescence *in situ* hybridization^87^ (MERFISH) coupled with immunostaining. We analyzed one GW23 (two technical replicates) and one GW14 (one technical replicate) sample. Following quality control, 340,114 cells were retained for downstream analysis. Across the tissue, we distinguished eleven transcriptomic clusters with differential spatial distributions that map onto the major cell types characterized in our multiome dataset. MGE and LGE progenitors included RG (*SOX2*+), IPC (*ASCL1*+) and dividing (MKI67+) populations. A neighboring cortical tRG-CRYAB cluster outside the GEs was also identified (**Fig. 4A, Supplemental Fig. 11, 12A-C, DataS3**). Neuronal populations included MGE-derived migratory interneurons, MGE-*CRABP1*+ cells, and developing striatal projection neurons, while non-neuronal lineages included microglia and vascular cells (**Fig. 4A**).

**Figure 4.**
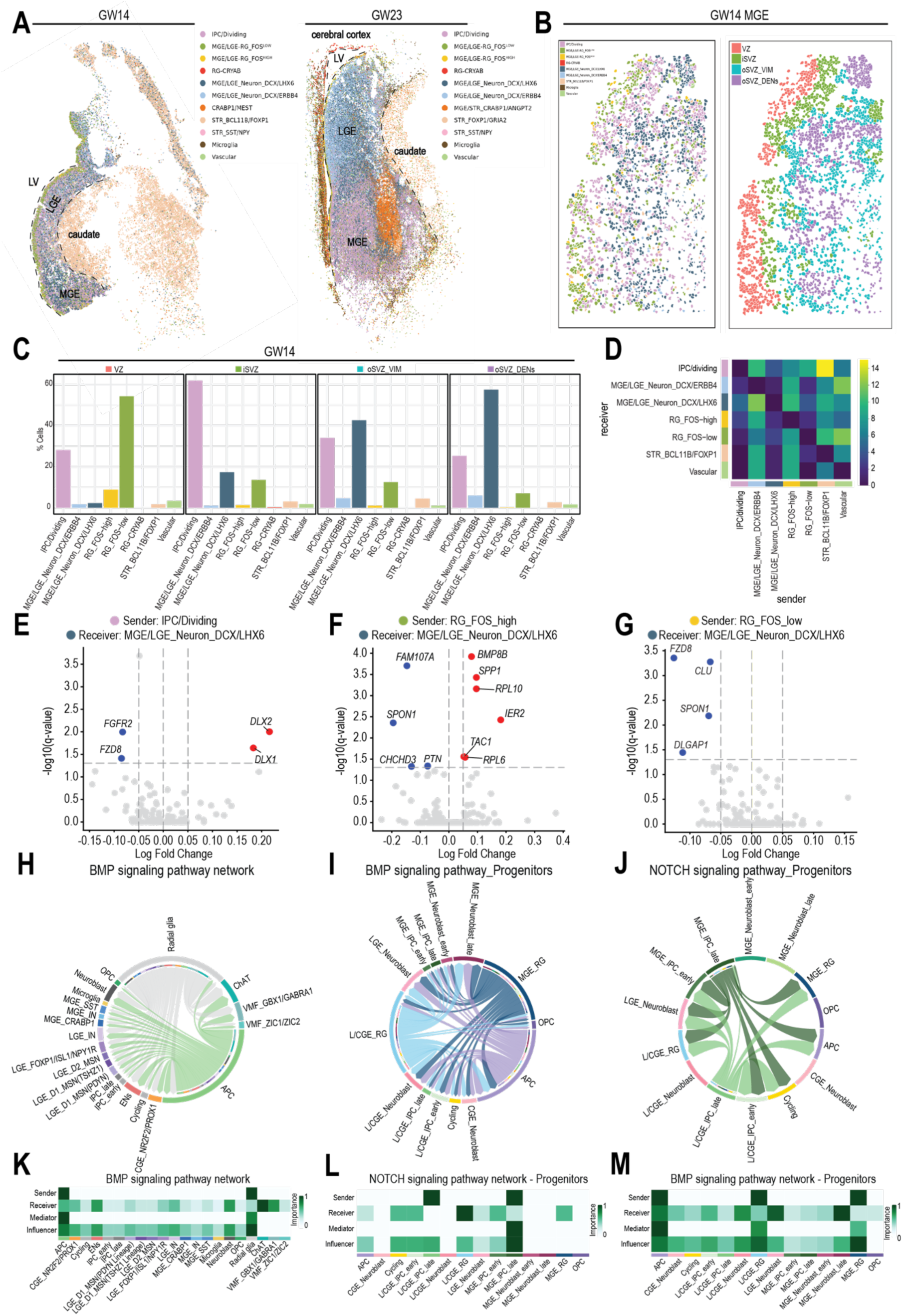
Spatial Transcriptomics of human MGE and LGE. **A**) Spatial transcriptomic analysis of the human LGE and MGE from GW14 and GW23. Cells are color coded by types. **B**) Zoomed in analysis of the GW14 MGE sample, cells are color coded by types or the niches to which they belong. **C**) The proportion of different cell types in individual niches. Niche numbers correspond to the legend in B. **D**) Type coupling heatmap showing sender-receiver interactions between cell types. Color intensity indicates the number of differentially expressed genes in receiver cells (y-axis) influenced by sender cells (x-axis) at FDR < 0.05. **E**, **F**, and **G**) Volcano plot of differentially expressed genes of MGE/LGE_Neuron_DCX/LHX6 in the neighborhood of IPC/dividing (**E),** RG_FOS_high (**F**), RG_FOS_low (**G**). **H**-**J**) Chord diagram showing cell-cell communication mediated by BMP signaling (**H**) in all cell types identified in the multiome data as shown in Figure1, BMP (**I**) and NOTCH (**J**) signaling in progenitor subsets as shown in Figure2. **K**, **L,** and **M**) Heatmap showing network centrality scores in the BMP signaling pathway network for in all cell types (**K**) identified in the multiome data as shown in Figure1, BMP (**L**) and NOTCH (**M**) signaling pathway network for progenitor subsets as shown in Figure2. Rows represent different signaling roles: Sender, Receiver, Mediator, and Influencer. Color intensity indicates the relative importance score for each role (scale: 0-1, light to dark green).

As a first step, we examined the distribution of progenitor cell populations. RG and IPC/dividing cells span the full extent of the MGE/LGE at both GW14 and GW23, indicating that these regions remain highly proliferative at these stages. (**Fig. 4A, B, Supplemental Fig. 11**). Within the RG compartment we observed two distinct cell states: a sparse, VZ-enriched RG_*FOS-high* population, scattered in SVZ, and a more abundant *RG_FOS-*low population marked by *PTPRZ1^+^*/*ATP1A2^+^*, distributed across both VZ and SVZ. Whereas our multiome analysis revealed regional transcriptional specialization among GE progenitors (**Fig. 2**), spatial transcriptomics uncovered an additional level of heterogeneity within the RG compartment defined by differential *FOS* activity. The two *FOS*-defined populations likely reflect distinct activation states^88,89^, rather than differences in RG subtypes or VZ–SVZ location.

Next, we focused on MGE-derived interneurons and identified subtypes destined for both cortex and striatum. Two main populations of migrating interneurons were identified as *DCX+/LHX6+* and *DCX+/ERBB4+*, the latter being more enriched in the migratory route towards LGE. We also observed striatal interneurons in the caudate marked by *SST* or *NPY* (**Fig. 4A, B**). Remarkably, our spatial maps revealed a previously unappreciated *CRABP1^+^*interneuron population within the oSVZ of the MGE at GW23 that has high *ANGPT2* expression and occupies a tightly packed, discrete territory. *CRABP1* expression was confirmed at the protein level through immunostaining; this population was negative for KI67, indicating that it is composed of postmitotic neurons (**Supplemental Fig. 13A, B).** From its discrete territory in the MGE oSVZ, the *CRABP1*⁺ population extends two streams: one toward the LGE, likely contributing to cortical interneurons, and another toward the striatum. with some already populating this region, suggesting it produces interneurons for both regions. This observation aligns with our multiome predictions that a subset of *CRABP1*⁺ cells may contribute to cortical interneurons, particularly of the PV fate. Sub-clustering of *CRABP1*⁺ cells and differential expression analysis revealed two populations: one enriched for LGE-directed cells expressing higher levels of *ANGPT2* and *MEST*, representing cortical contributions, and the other primarily containing cells enriched in the striatum expressing higher levels of *CRABP1* and *ETV1* (**Supplemental Fig. 12D-F**). Our findings extended previous observations of species-dependent differences in the ultimate destination of *CRABP1*⁺ interneurons^21,90^ by revealing a discrete *CRABP1*⁺/*ANGPT2*⁺ domain in the human MGE at GW23 that contributes to both cortical and striatal interneurons.

### The human MGE is divided into distinct cellular subdomains

We next focused on the putative primate-specific cytoarchitecture^30,31^ of the MGE. Using the GW14 MGE sample, we jointly interrogated cell type identities and their spatial distribution by performing spatial domain analysis based on k-nearest neighbor graphs^91^. We identified four distinct spatial domains characterized by different relative cell type proportions along the three proliferative compartments: VZ, iSVZ, and oSVZ^31^ (**Fig. 4C**). The VZ compartment (**Fig. 4B**, red) was enriched for RG, whereas the adjacent iSVZ (**Fig. 4B**, green) was predominantly composed of *ASCL1^+^* IPCs, consistent with prior reports^31^ and our immunostaining data (**Fig. 4C; Supplemental Fig. 13D-E**). Notably, the expanded oSVZ compartment further resolved into two distinct domains (**Fig. 4B**, teal and purple,): one highly enriched for *DCX+/LHX6+* cells corresponding to DENs (oSVZ_DENs), and the other comprising a vimentin+ progenitor cluster (oSVZ_VIM) with mixed *ASCL1^+^*and *DCX^+^* cells organized around DENs (**Fig. 4C; Supplemental Fig. 13D-E**).

To further characterize the three-dimensional organization of the human MGE, we employed electron microscopy to analyze 1.5 µm serial semithin sections of DENs at GW23 (**Fig. 5A-D,)**. From the VZ to the inner portion of the oSVZ, we observed that multiple DENs are often “walled off” by radial glial fibers that enclose them within a shared structural unit (**Fig. 5B)**. Examination of DENs within these structural units revealed that they are not spherical but instead form columnar structures that interconnect to form a trabecular-like architecture (**Fig. 5B-D)**. Within each column, IPCs with occasional RGs separate individual DENs along the apical-basal axis, as shown by ASCL1 immunostaining (**Fig. 5E, Supplemental Fig. 13D**) and consistent with the high proportion of IPCs observed in our spatial analysis (**Fig. 4A-C**). By integrating progenitor immunostaining, ultrastructural analyses, and spatial transcriptomics, we find that DENs are organized along a periphery-to-center differentiation gradient. Radial glial cell bodies and fibers define an outer shell, followed by IPC-enriched zones containing scattered RG. Toward the center, IPCs progressively transition to neuroblasts, culminating in a neuroblast-rich core (**Fig. 5D-F**). This organization defines DENs as structured developmental units that may support coordinated proliferation, migration, and lineage progression within the MGE.

**Figure 5.**
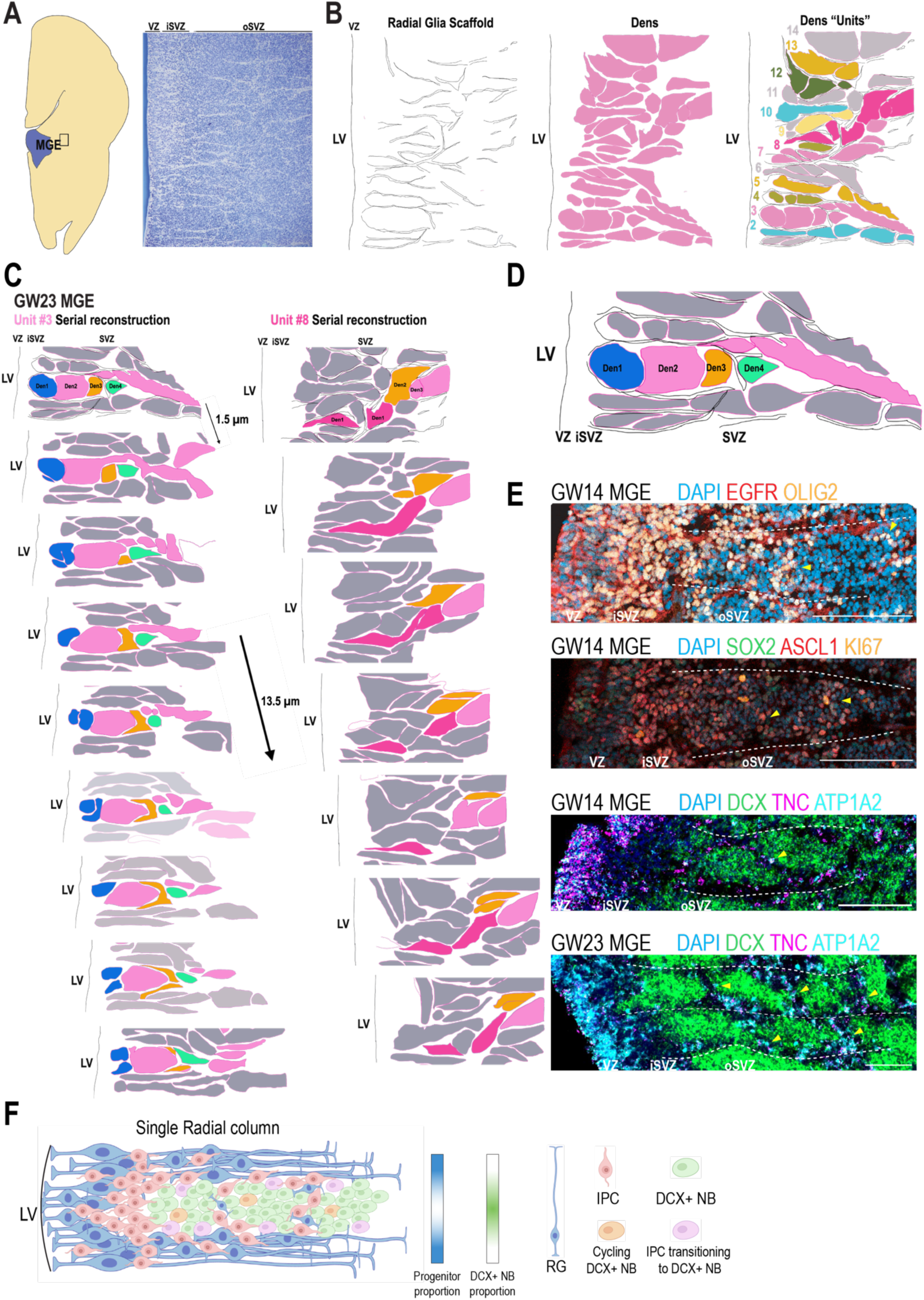
**Integrated spatial analysis and structural reconstruction of the MGE cytoarchitecture**. **A**) Nissl-stained section of the human MGE at GW23. **B**) Schematic representations of the reconstructed radial glial scaffold, DENs, and DEN “units” of the human MGE at GW23. **C**) Serial EM–based reconstruction of DEN structure. **D**) Schematic of a reconstructed DEN radial column highlighting columnar organization. **E**) Immunostaining and RNAscope demonstrating the spatial organization of progenitor populations and DENs within a single radial column. **F**) Schematic summarizing cell-type distributions within a single radial column in the human MGE.

### Cell-cell communication in the developing human MGE shows inter-progenitor signaling

Our spatial transcriptomics and electron microscopy analyses reveal a nuanced cellular organization of human MGE DENs, that likely arises from the interplay of intrinsic and extrinsic signaling. To probe regulatory mechanisms underlying DEN organization and cellular dynamics, we applied node-centric expression modeling to the GW14 spatial dataset^92^, uncovering cell type–specific interaction networks. (**Fig. 4D-G**). This analysis predicted that progenitor cells modulate the *DCX+/LHX6+* neuroblast transcriptional program: *ASCL1+*intermediate progenitor cells (IPCs) positively regulated *DLX1* and *DLX2*, while repressing *FZD8*, *FGFR2* (Wald test; FDR-correct p-value < 0.05) and *NOTCH1* (FDR-correct p-value < 0.10). In parallel, RG were predicted to negatively regulate *FZD8* expression (FDR-correct p-value < 0.05) (**Fig. 4E-G)**. These coordinated regulated shifts point to a shared trajectory characterized by attenuation of progenitor programs and adoption of a neuroblast identity. To supplement our spatial analysis and identify additional signaling mechanisms operating within the developing MGE, we performed ligand–receptor analysis using CellChat^93^ on our multiome dataset (**Fig. 4H-M**). We identified BMP signaling from RG to both NB (RG-NB) and MGE-derived interneurons (RG-MGE-IN), as well as NOTCH signaling from IPCs to NB (IPC-NB) and RG. These interactions suggest that BMP and NOTCH pathways may serve as mediators of progenitor-NB communication^94^. These findings support a model in which progenitor populations orchestrate DEN cellular dynamics by regulating the balance between proliferation and differentiation.

### MGE-enriched GE organoids replicate human MGE subtypes and GE-oRG

Mechanistic investigation of DEN formation has been challenging due to limited access to human primary tissue, the rapid disorganization of *ex vivo* slice cultures, and the lack of an organoid model that reproduces DEN structures. To overcome this, we generated MGE-enriched GE organoids that recapitulate key cell types, reliably form DEN-like structures, and are compatible with genetic manipulation and live imaging. We adapted a published 2D MGE protocol^95,96^ for organoid culture (**Fig. 6A**). Immunostaining confirmed MGE regional identity with 66% of cells at DIV21 expressing the MGE-specific marker, NKX2.1^7^ (**Fig. 6B, C**). Single cell RNAseq of GE organoids at DIV51 and DIV164 further confirmed cellular diversity and regional identity, revealing progenitors (*PTPRZ1*, *VIM*, *ATP1A2*, *SOX2*) and neuron clusters (*DCX*) derived from all three GE sources; MGE (*LHX6*, *SST*, *CRABP1*), LGE (*SP8*, *FOXP1*) and CGE (*SCGN*, *PROX1*) (**Fig. 6D**). To examine the correspondence between organoid-derived progenitor cells with a focus on GE-oRG and their native counterparts, we performed live imaging of organotypic slices prepared from organoids (derived from four independent cell lines) at DIV25 and DIV47 when large numbers of KI67+ dividing cells are present (**Fig. 6E-G; Supplemental Fig. 14A**). Focusing on GE-oRG cells (**Fig. 3**), MST was observed in 44.0% of cell divisions at DIV25 and 42.1% at DIV47 (**Fig. 6E; Supplemental Fig. 14B**). The mean MST translocation length was 43.3mm at DIV21 and 45.0mm at DIV47 (**Fig. 6F**), closely resembling values reported for GE-oRG in primary MGE (46.3 µm) and LGE (47.8 µm) tissue (**Fig. 6G**). Consistent with these live imaging results, snRNA-seq confirmed the presence of GE-oRG markers (*ATP1A2, PTPRZ1,* and *HOPX*; **Fig. 6D).** Collectively, these findings indicate that GE organoids contain major cell types found in the GEs, including GE-oRG, with enrichment of MGE subtypes supporting their use as a high-fidelity model to study ganglionic eminence development.

**Figure 6.**
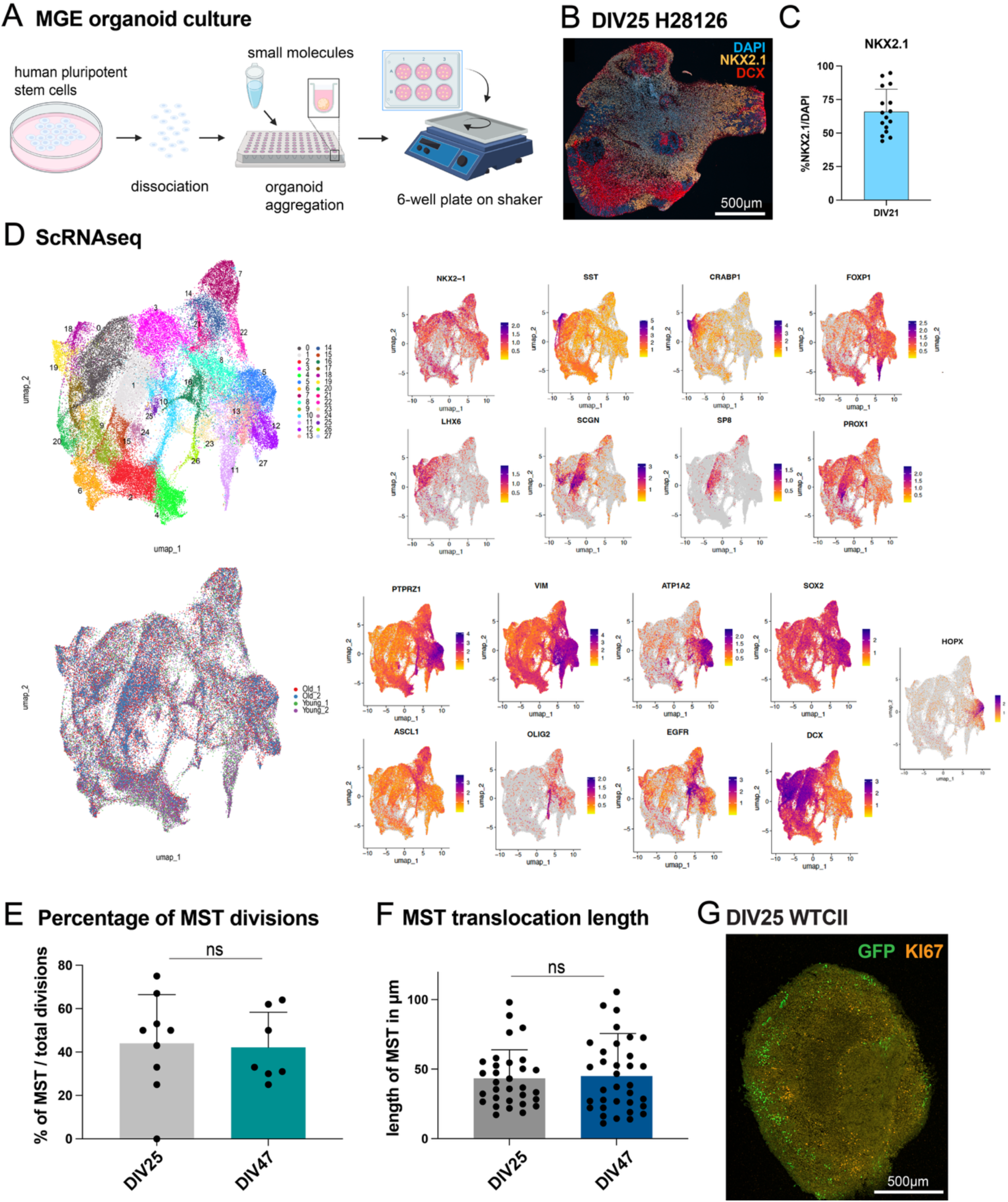
GE organoids show enrichment of MGE cell types and replicate cell types of the human MGE. **A)** Experimental scheme of organoid culture. **B)** Example immunostaining of H28126 line at DIV25. Scale bar is 500mm. **C)** Quantification using Imaris software of NKX2.1 protein expression at DIV21 (3 cell lines). **D)** ScRNA-seq analysis of MGE organoids with UMAP clustering revealing 29 distinct cell clusters based on principal component analysis (PCA). **E)** Proportion of MST divisions relative to non-MST divisions in organoids at DIV25 and DIV47 (3 cell lines). **F)** Average MST translocation length. **G)** Immunostaining of organoid slice following live imaging and infection with Ad-CMV-GFP.

### PCDH19 regulates DEN formation in human MGE organoids

We next examined DEN formation in our MGE-enriched GE organoids. At DIV70–90, immunostaining showed the emergence of pseudo-DENs-like structures with mutually exclusive DCX and VIM expression but inverted organization with VIM+ clusters surrounded by DCX+ cells (**Fig. 7A)**. At DIV150, DENs were surrounded by VIM+ fibers with sparser DAPI staining, more closely resembling those in primary tissue, that we refer to as mature DENs (**Fig. 7C**). These structures were observed across multiple cell lines and colocalized with LHX6 staining indicating MGE identity (**Supplemental Fig. 14C-E**). Two types of DENs and surrounding progenitor clusters have been described in the human MGE: type 1 progenitor-adjacent DENs are well-rounded, densely packed and closer to the iSVZ, while type 2 progenitor clusters and DENs are thin, elongated streaks within the oSVZ^31^. Interestingly, we identify DENs with morphologies similar to both subtypes in our organoids at DIV150 (**Fig. 7C**). Using RNAscope, we also observe that *ATP1A2*- and *TNC*-expressing RG surround *DCX*+ DEN structures as observed in primary tissue (**Fig. 7B; Supplemental Fig. 6C, D**). Dissociated organoids transplanted into mouse cortex at DIV130 and harvested 7 weeks after transplantation were able to form mutually exclusive clusters of DCX or VIM expressing cells (**Fig. 7D**) similar to dissociated human MGE^30^, highlighting that DEN formation is influenced by intrinsic signals.

**Figure 7.**
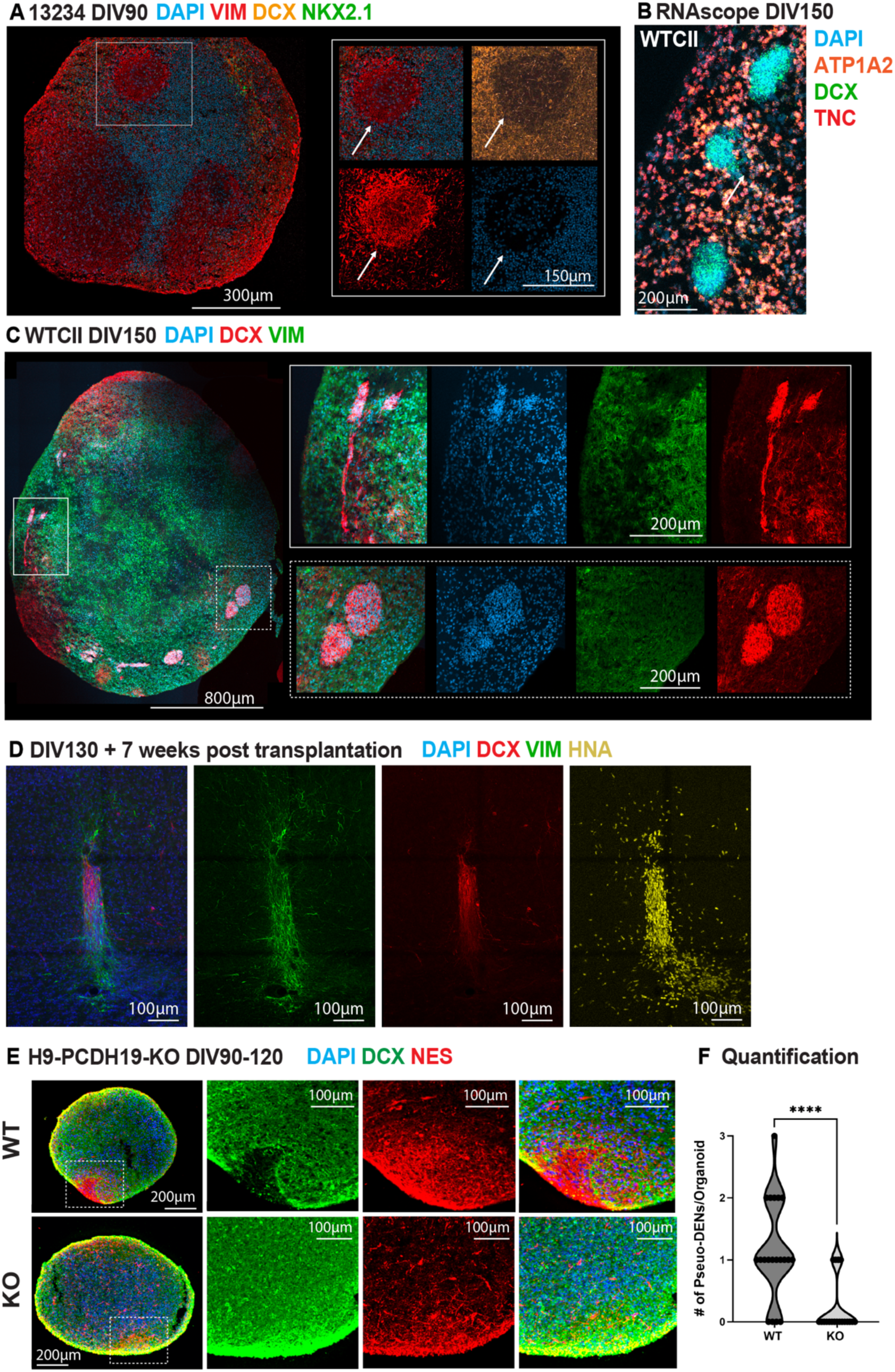
PCDH19 regulates DEN formation in MGE-enriched GE organoids. **A)** Immunostaining at DIV90 showing immature DENs, characterized by mutually exclusive expression of VIM and DCX (white arrow), without VIM fibers surrounding DCX⁺ clusters. **B)** RNAscope of organoids at DIV150 shows expression of *ATP1A2+/TNC+* RG surrounding *DCX+* DENs, replicating primary MGE. **C)** Immunostaining at DIV150 showing mature DENs with VIM-rich processes enveloping DENs. DENs (dashed white line) resembling type1 and type 2 (solid white line) progenitor surrounded DENs are highlighted. **D)** Immunostaining of transplants of dissociated GE organoids that were transplanted into mouse cortex at DIV130 (organoid age at day 0 of transplantation) and harvested 7 weeks after transplantation. **E)** Representative images of immunostaining for doublecortin (DCX; green) and Nestin (red) in 4-month MGEOs derived from *PCDH19* CRISPR knockout (KO; bottom row) and wildtype isogenic control (WT; top row) hPSCs. Nuclei are shown in blue with bisbenzimide staining. **F)** Quantification of DCX-negative pockets in 4-month immunostained organoids of two cell lines shows a statistically significant decrease in DCX-regions in *PCDH19* KO MGEOs compared to WT MGEOs (Student T-test ****p=0.0001). Total of 4 independent differentiations (diffs) (n=12-13 organoids from n=3 female diffs, and n=7 organoids from 1 male diff).

How DENs structures are formed remains unclear, but differential expression of the protocadherins PCDH10 and PCDH19 has been proposed^30^. Given that PCDH19 is enriched in DENs, we investigated its role using MGE organoids^97^ generated from two *PCDH19* KO lines and isogenic controls^97,98^. We observed a significant reduction in the number of pseudo-DENs in *PCDH19* KO lines compared to controls (p**=0.0018) (**Fig. 7E, F)**, suggesting that PCDH19 may be involved in the regulation of DEN formation in the human MGE. In sum, human MGE-enriched GE organoids faithfully recapitulate primate MGE cytoarchitecture, including progenitor clusters and DENs. Loss of *PCDH19* disrupts their formation, highlighting its essential role and offering a tractable system for mechanistic studies of interneuron niche assembly.

## Discussion

Our integrated analysis of the developing human GE combines single-nucleus transcriptomics, chromatin accessibility profiling, spatial transcriptomics, live-cell imaging, electron microscopy, and organoid models to provide a comprehensive framework for understanding progenitor diversity across the developing ganglionic eminences, as well as insights into spatial organization and mechanistic formation of DENs.

### GE-oRG may drive GE expansion during mid-gestation

Previous studies have identified non-epithelial progenitors in the ganglionic eminences^21,30,31^. Here we characterize a population of outer radial glia-like progenitor cells in the SVZ and oSVZ of the human MGE, LGE, and CGE that we term GE-oRGs. We propose that GE-oRGs are defined by a constellation of features that collectively distinguish them from both cortical oRGs and GE ventricular radial glia. First, similar to cortical oRGs, GE-oRGs reside within the SVZ/oSVZ. Second, transcriptional divergence between vRGs and oRGs is less pronounced in the ganglionic eminences than in cortex^78^. Third, GE-oRGs undergo MST, but with region-specific dynamics: translocation distances are shorter in the MGE and LGE and longer in the CGE compared to cortical oRGs. Fourth, GE-oRGs have a unipolar morphology.

The molecular profiles of GE progenitor subtypes likely reflect a continuous gradient of gene expression rather than discrete segregated cell states. This organization is similar to the apical (B1 cells) and non-apical (B2 cells) neural stem cells in the adult V–SVZ that share overlapping transcriptional and morphological features but B1 cells are located in the VZ and B2 cells in the SVZ^99^. The human MGE increases in volume from approximately 115 mm³ at GW 18 to 158 mm³ at GW 22^30^. Given that dividing GE-oRGs account for a substantial proportion of mitotic events, we propose they may contribute to progenitor zone expansion in the ganglionic eminences, analogous to the role of cortical oRGs in driving cortical progenitor zone expansion^79,100,101^.

### Convergence and divergence of dorsal and ventral RG

Our comparison of cortical and ganglionic eminence RG revealed that regional differences are more pronounced at the epigenomic than the transcriptomic level. Overlap of marker chromatin peaks is limited (19.3% of marker peaks versus 50.2% of marker genes, **Supplemental Fig. 9D, 8D**), and this divergence extends beyond marker loci to the broader regulatory landscape. Only 53.7% of highly variable chromatin peaks are shared between regions, compared with 77.7% of highly variable genes (**Supplemental Fig. 9D, 8D**). These findings indicate systematic epigenomic divergence that is not captured by transcriptional profiles alone. Indeed, enhancers are inherently more cell-type specific than transcriptional signatures alone^102,103^. The pronounced epigenomic differences likely arise from two complementary mechanisms. First, region-specific developmental niches impose distinct regulatory architectures that configure chromatin accessibility landscapes^39,104,105^. Second, epigenetic priming establishes enhancer accessibility patterns that reflects progenitor competence in advance of downstream transcriptional programs^40,86^. This priming architecture reveals a hierarchical regulatory framework in which chromatin accessibility defines developmental potential more sharply than gene expression, pre-positioning enhancers to activate lineage-specific transcriptional programs that emerge as RG differentiate.

### The diversity of primate GE progenitors may have contributed to the evolutionary expansion of interneuron populations

The human ganglionic eminences undergo profound evolutionary expansion in both cellular composition and organizational complexity, supporting the increased production of GABAergic interneurons required for the enlarged primate forebrain^30,31,106^. MGE-derived striatal interneurons such as TAC3+ cells^29,90,107,108^ and cortical GABAergic interneurons including CGE-derived populations^108–110^ are present at higher proportions in primates than in rodents. This expansion has been proposed to arise from a multi-tiered architecture of GE progenitor diversity and organization unique to primates ^21,25,30,31,34^. We propose that neuronal amplification may be driven by three hierarchically organized progenitor tiers: (1) non-apical GE outer radial glia (GE-oRGs), (2) intermediate progenitor cells (IPCs) and (3) proliferative neuroblasts. First, the broad distribution of *ATP1A2+/TNC+/PTPRZ1+* RG throughout the oSVZ of the human LGE and MGE, especially encircling MGE DENs, combined with their high frequency of MST, suggests that GE-oRGs contribute substantially to progenitor amplification. Second, consistent with previous observations of expanded progenitor populations in the primate MGE^21^, we find that ASCL1⁺ IPCs form a dense inner SVZ layer in human tissue, a structure absent in the mouse^111^. Ultrastructural and immunohistochemical analyses further show that ASCL1⁺ IPCs separate adjacent MGE DENs, positioning them to efficiently supply newly generated neuroblasts to DENs. Third, DENs contain DCX+/KI67+ proliferative neuroblasts that retain proliferative capacity despite expressing neuronal markers^30^. Together, the expansion of GE-oRGs, IPCs within the iSVZ and around DENs, and abundant proliferative neuroblasts within DENs provide a cellular framework for primate-specific amplification of interneuron production.

### A segregated CRABP1⁺/ANGPT2⁺ MGE domain may underlie primate interneuron diversity through niche-driven specification

Spatial transcriptomics revealed that by GW23 *CRABP1⁺/ANGPT2*⁺ interneuron precursors consolidate into a well-defined territorial domain within the MGE. Although CRABP1⁺ cells are detectable at earlier gestational ages, they form a spatially organized territory within the MGE oSVZ at GW23 that may facilitate the generation of interneuron diversity destined for both cortex and striatum. The CRABP1⁺/ANGPT2⁺ domain comprises postmitotic neurons and gives rise to two distinct migratory streams, one extending toward the LGE and cortex and another toward the striatum. This bifurcation may be supported by niche-specific molecular programs that differentially modulate migratory cues. Persistent *NKX2-1* expression, known to direct striatal interneuron migration^112^, could bias a subset of cells toward striatal compartments via its downstream transcriptional network. Concurrently, ANGPT2 secretion by these cells may promote endothelial–neural crosstalk^113,114^, providing metabolic, growth factor, and angiogenic signals, to stabilize the niche and support maturation. Together, these findings identify the spatially confined CRABP1⁺/ANGPT2⁺ MGE domain as a specialized developmental hub that integrates positional identity, migratory routing, and niche signaling to guide the development of primate-expanded interneuron subtypes.

### IPC and oRG promote neuronal amplification within columns of MGE DENs

Unlike the developing neocortex, the MGE does not show laminar organization^115,116^. Instead, newborn GABAergic interneurons in the MGE organize into neuroblast clusters (DENs) that are surrounded by spatially restricted progenitor clusters^30,31^. This structure is absent in mice, and although MGE DCX+ clusters are observed in pig MGE, the associated progenitors are broadly dispersed^30,31,33^. These comparisons suggest that true DENs surrounded by restricted progenitor clusters may be a primate-specific organizational feature. Serial EM reconstruction shows that RG processes form columns of interconnected fibers that surround and separate multiple stacked DENs. While radial glial processes provide the structural scaffold organizing DEN architecture, the DCX+/KI67+ neuroblasts within DENs have been proposed to serve as a major amplifying cell type in the MGE^30^. Our structural analysis and live imaging extend this model by demonstrating that highly proliferative GE-oRGs and IPCs that surround and separate individual DENs are additional drivers of neuronal amplification within the human MGE. Moreover, our analysis of cell-cell communication further supports this model. BMP-mediated signaling from RG to neuroblasts, and NOTCH-mediated signaling from IPC to neuroblasts, suggest that GE-oRGs and IPCs may actively sustain DEN growth by continually supplying newly generated neuroblasts. The trabecular organization of DENs thus emerges as a multicellular scaffold that integrates structural support, proliferative amplification, and local signaling, to support the unique migratory behaviors and developmental trajectories of GABAergic neurons.

### MGE organoids pinpoint the role of PCDH19 in DEN formation

Previous studies have described MGE organoids with reliable cellular composition^98,117–120^, including one study reporting DCX+ cell clusters^121^. However, to our knowledge type 1 and 2 progenitor clusters^30,31^ and mature DEN-like structures have not been observed in organoids before our study. Differential protocadherin expression has been proposed to play a role in DEN formation^30^. Using MGE organoids derived from *PCDH19* knockout (KO) PSCs, we show that *PCDH19* is required for DEN formation. Loss-of-function mutations in *PCDH19* cause Protocadherin-19 Clustering Epilepsy (PCE), a severe early-onset form of epilepsy^122–127^ whose prevailing pathogenic mechanism involves abnormal cell sorting driven by differential adhesion between mutant and wild-type cells. This aberrant segregation has been demonstrated in both mouse models^128^ and human cortical organoids^98,129^. In mosaic cortical organoids containing a mixture of *PCDH19*-knockout and wild-type cells, the two populations spontaneously segregate into separate domains, a process termed cellular interference^129^. Notably, mosaicism is required for this segregation to occur^129,130^. Extending this framework to the developing MGE, we propose that a differential protocadherin combinatorial code likely underlies DENs development, in which PCDH19-expressing neuroblasts segregate apart from PCDH10-expressing early progenitors, thereby driving the self-organization of DENs. This mechanism provides a unifying explanation linking protocadherin-mediated cell sorting to primate-specific progenitor niche architecture and to the pathogenesis of PCDH19-associated epilepsy.

## Supporting information

DataS4

DataS3

DataS2

DataS1

## Acknowledgements

The research for this study received financial assistance from a PhD fellowship to Clara Siebert from the Boehringer Ingelheim Fonds (BIF) and was also supported by NIH grants 1R35NS137344, 1UM1MH130991 to Arnold Kriegstein. The human fetal material was provided by the Human Developmental Biology Resource (HDBR) (http://hdbr.org). This publication was supported by and coordinated through the Brain Initiative Cell Atlas Network (BICAN). We thank Alex Pollen for sharing the NKX2.1-GFP reporter cell line. We thank John Rubenstein for helpful discussions.

## SUPPLEMENTARY FIGURES

**Supplementary Figure 1.**
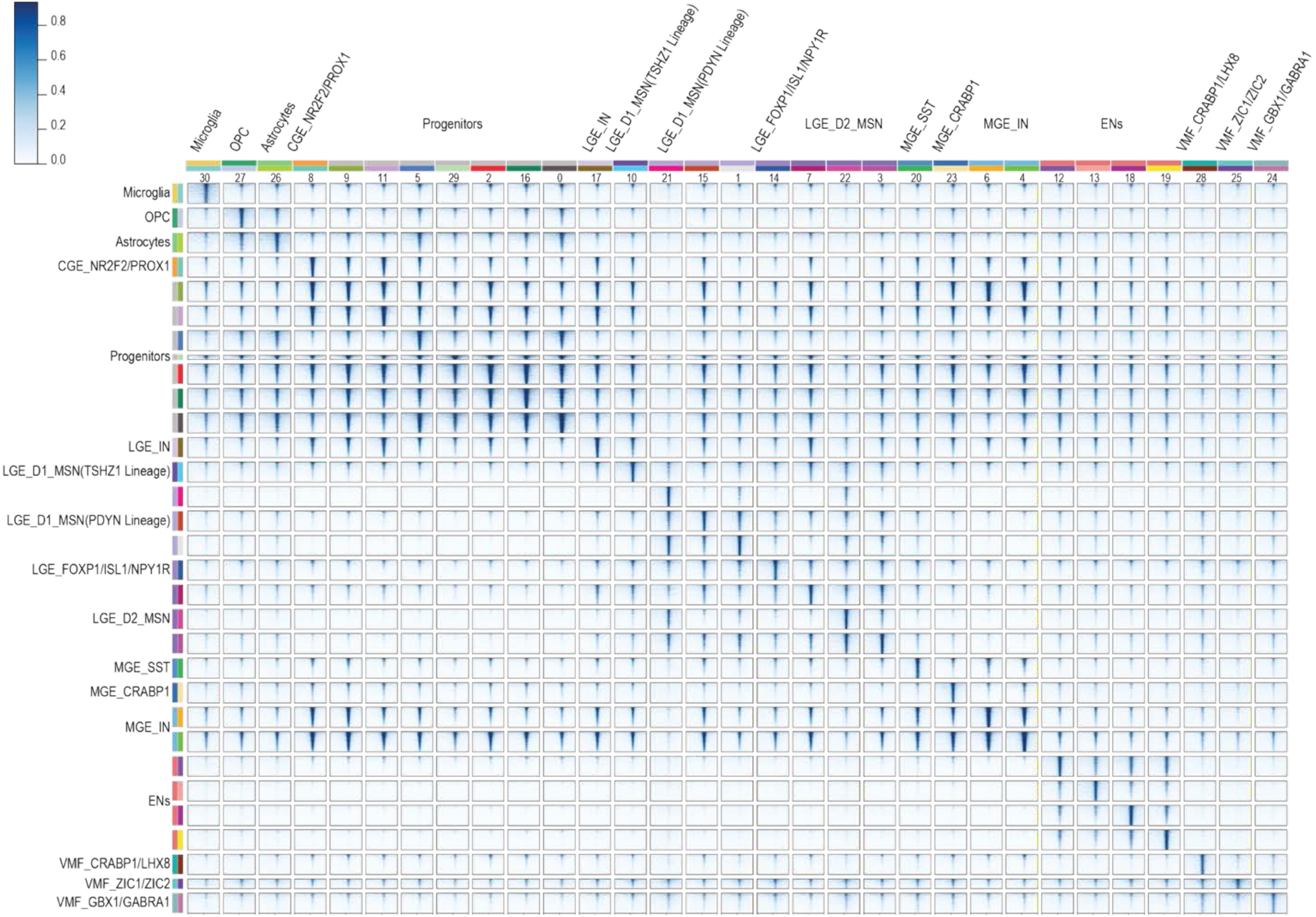
Cell-type–specific open chromatin profiles. A heatmap visualizing the top 1,000 DARs per cell type. For each peak, blue intensity corresponds to signal strength within a ±3 kb window around the peak center.

**Supplementary Figure 2.**
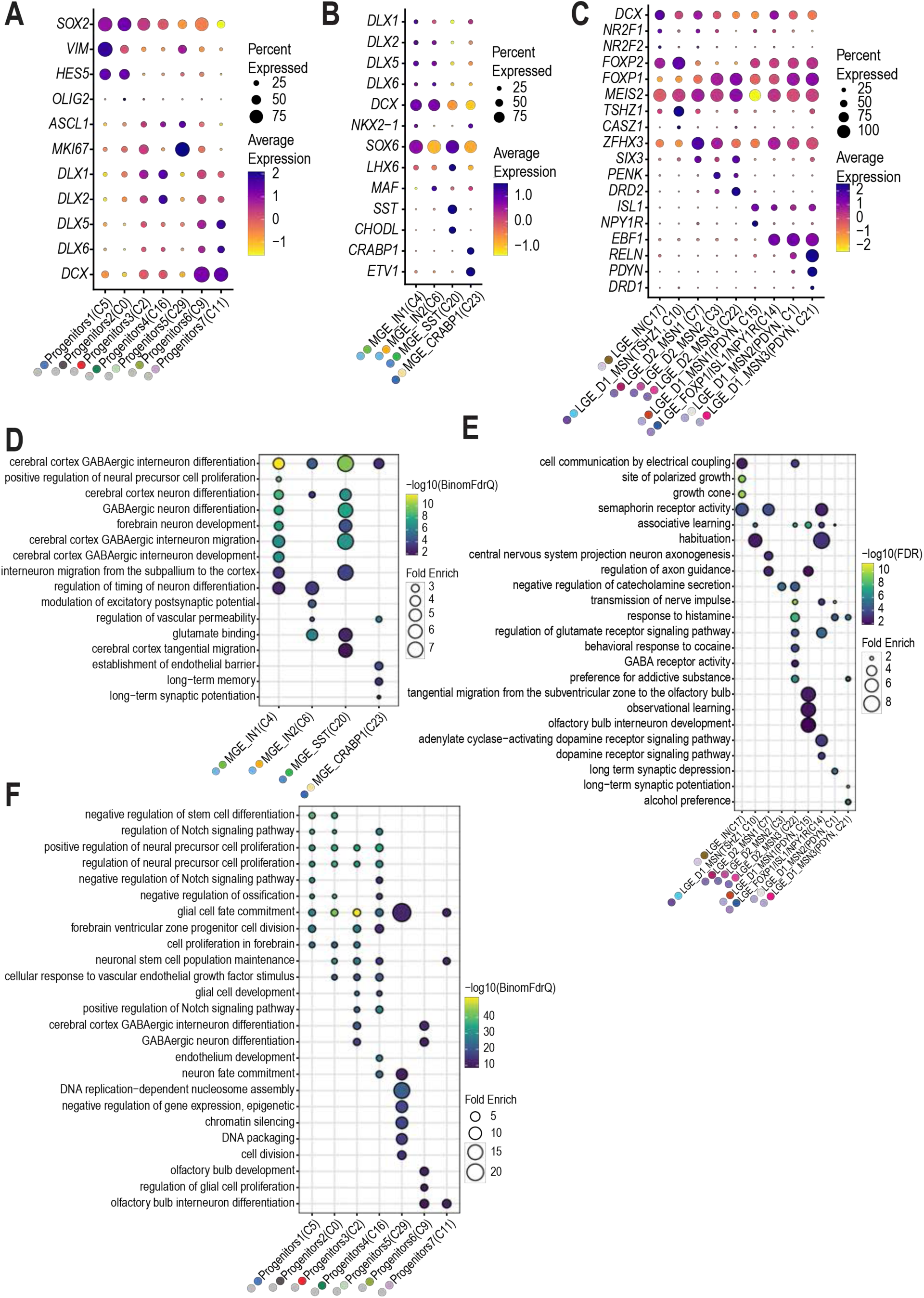
Transcriptomic and Epigenomic Signatures Across GE Lineage. **A–C)** Dot plots showing the mean expression levels (color) and the percentage of cells expressing each marker gene (dot size) across progenitor (**A**), MGE-lineage (**B**), and LGE-lineage clusters (**C**). **D-F**) Dot plots showing the most enriched terms obtained from DARs in progenitor (**D**) MGE-lineage (**E**), and LGE-lineage clusters (**F**) using GREAT analysis.

**Supplementary Figure 3.**
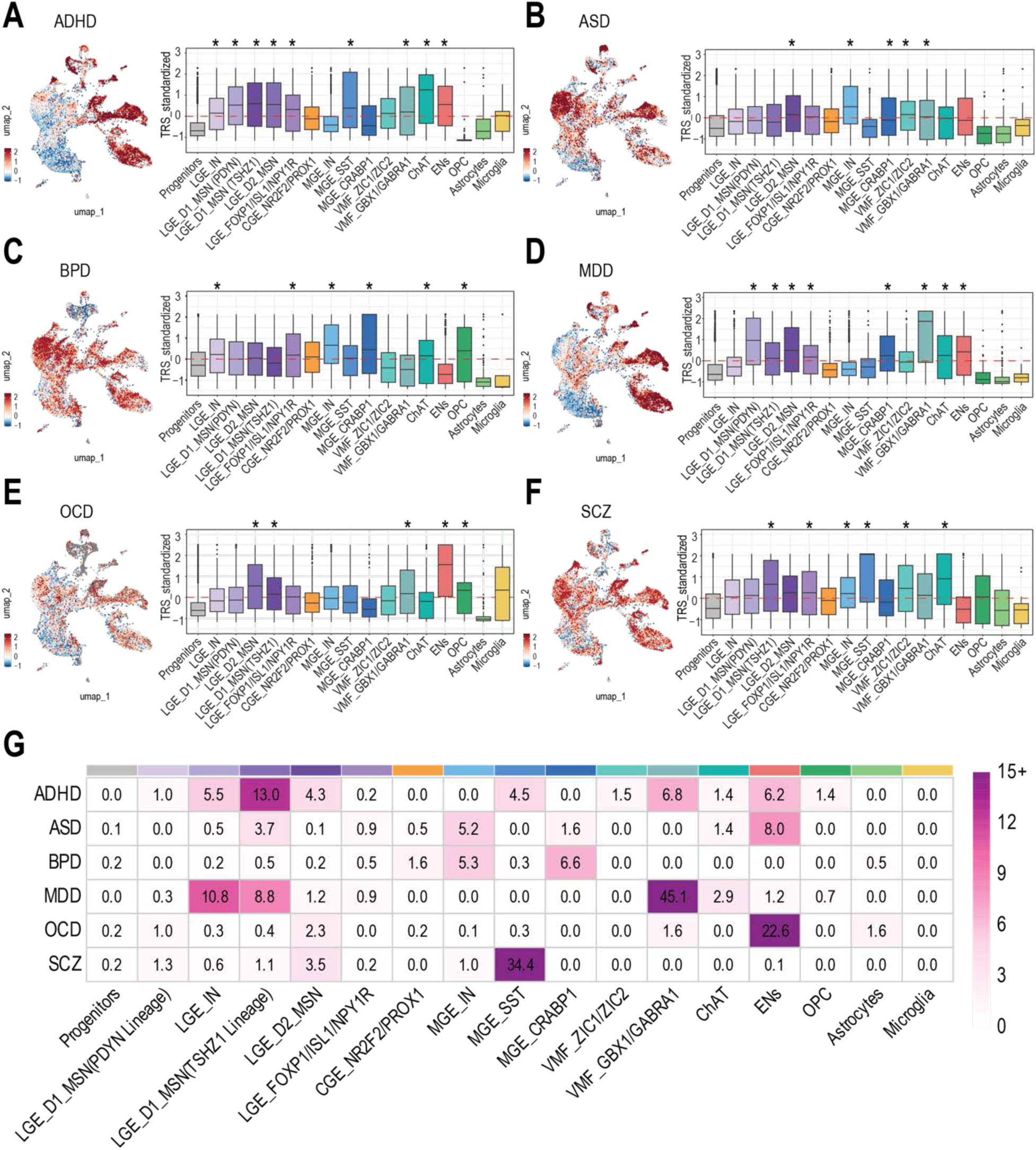
Cell-type association with human cognition and brain disorders. **A, B, C, D, E, and F**) Standardized per-cell SCAVENGE TRS for six brain disorders, including ADHD (A), ASD (B), BPD (C), MDD (D), OCD (E), and SCZ (F). **G**) The proportion of cells with enriched trait relevance across cell types.

**Supplementary Figure 4.**
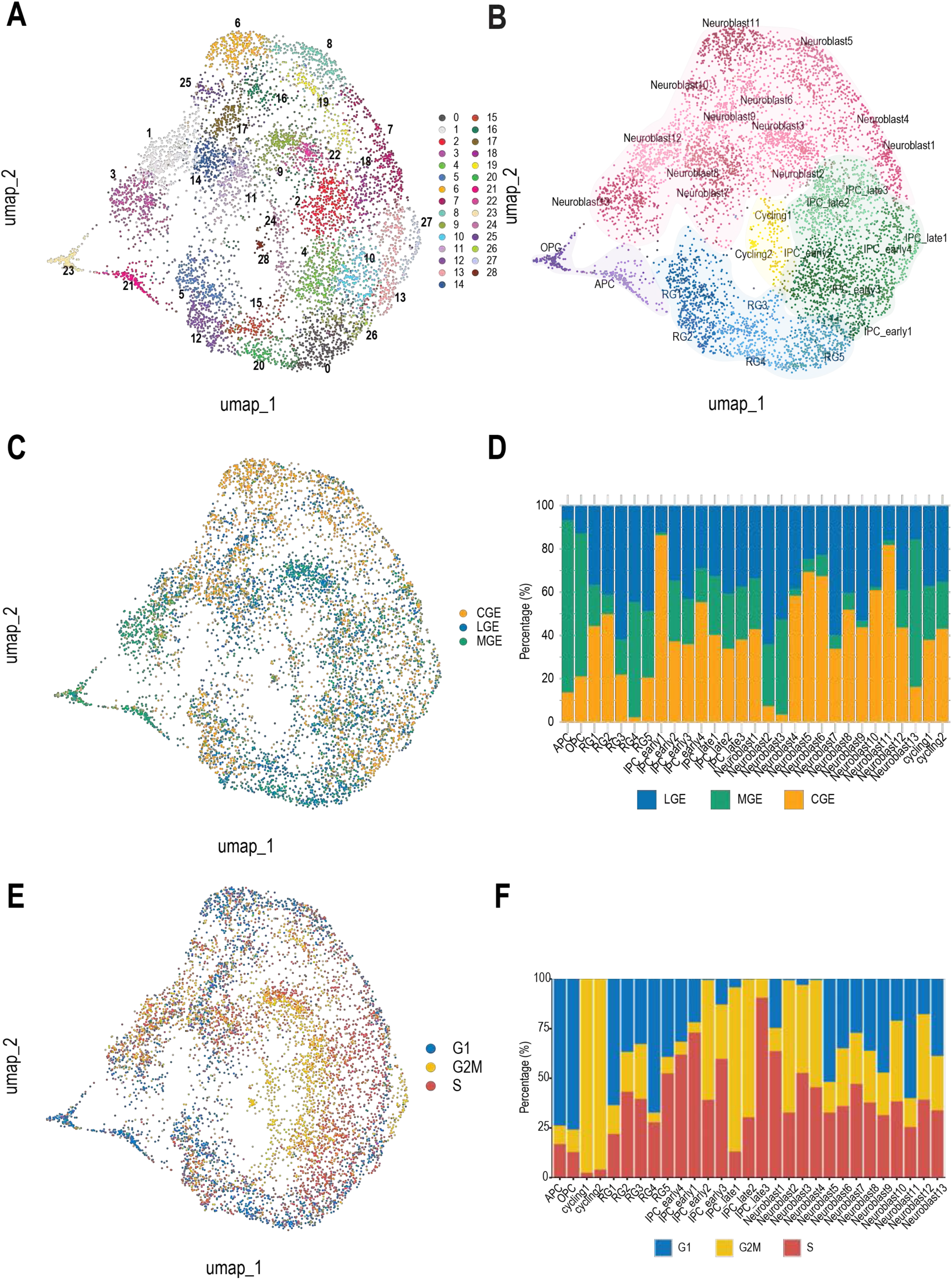
Landscape of Progenitor Diversity in the developing human GE. **A-C**) UMAP visualization of progenitors from merged GW14 and GW20 LGE, MGE, and CGE datasets, colored by cluster (**A**) or cell type (**B**) or regional origin based on dissection (**C**). **B**) Bar plot showing the regional composition (LGE, MGE, CGE) of each progenitor cell type. **E**) UMAP visualization of the same progenitor populations colored by cell cycle phase. **F**) Bar plot showing the distribution of cell cycle phases across progenitor cell types.

**Supplementary Figure 5.**
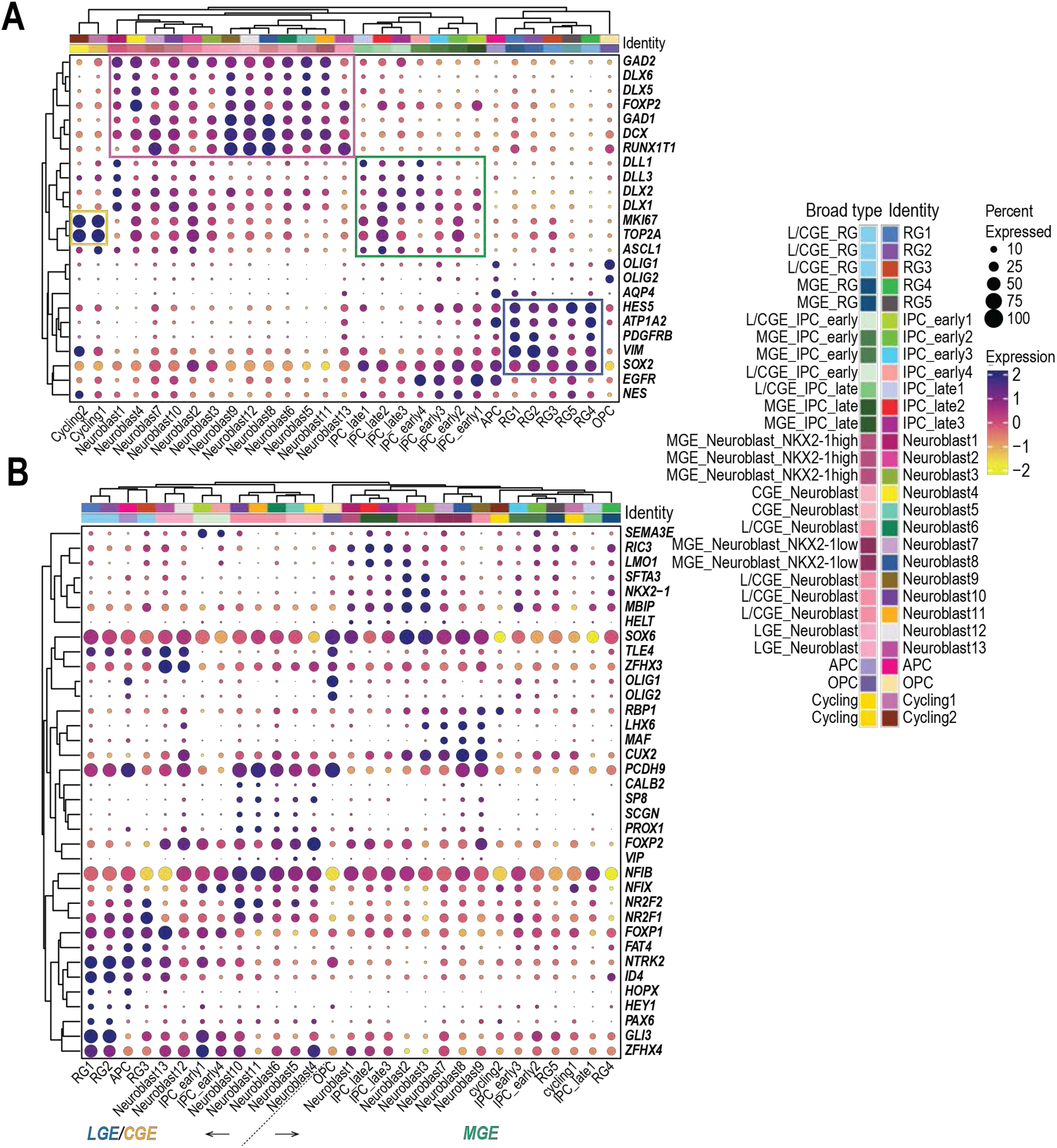
Landscape of Progenitor Diversity in the developing human GE. **A**, **B**) Clustered dot plots showing expression of progenitor marker genes (**A**) and regional marker genes (**B**) across all progenitor populations.

**Supplementary Figure 6.**
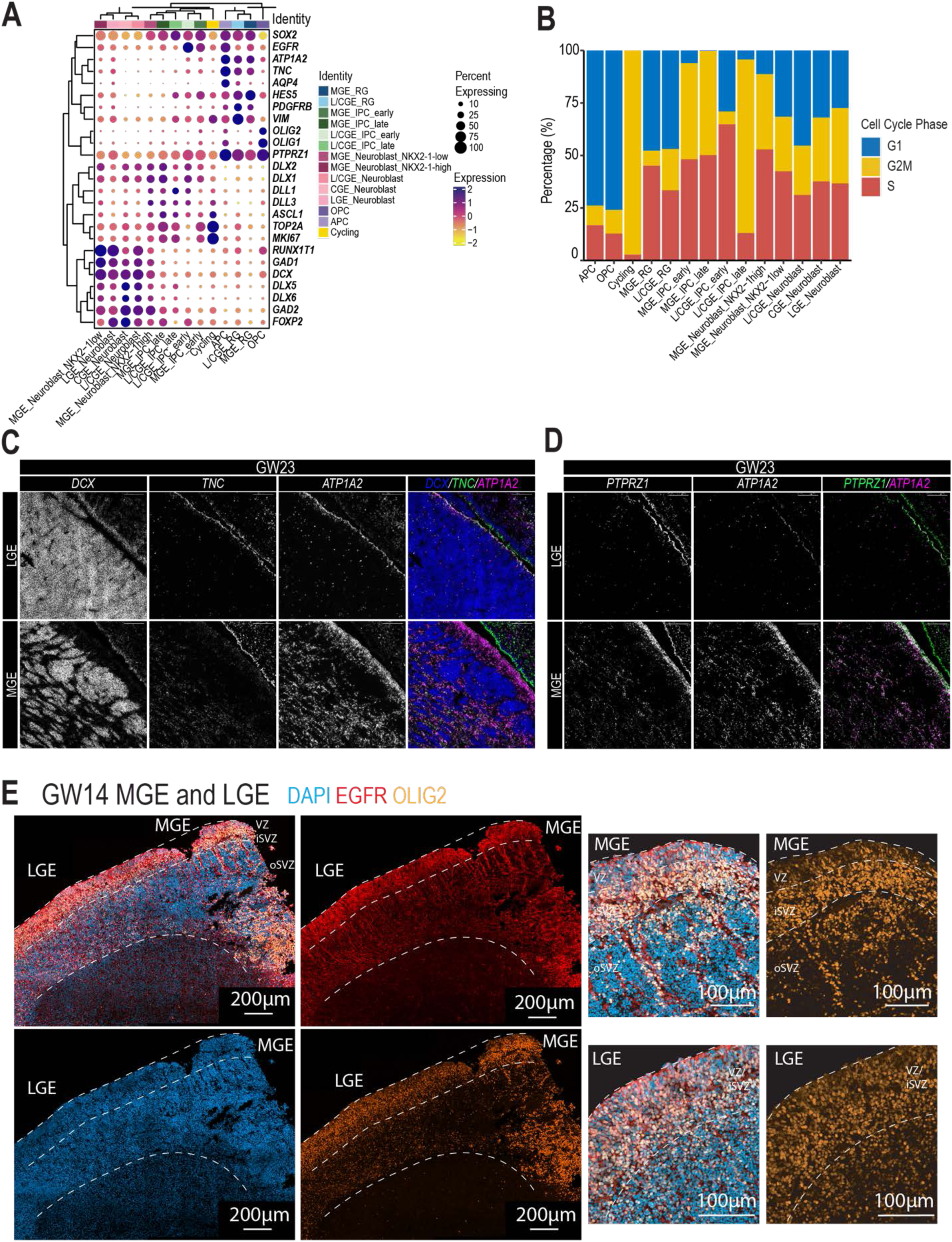
Molecular and Cellular Characterization of GE Progenitors. **A**) Clustered dot plot of progenitor marker gene expression across broad progenitor types. **B**) Bar plot showing the proportion of cell cycle phase across broad progenitor types, with bars color-coded by S, G1, and G2 phases. **C**, **D**) RNAscope showing expression of RG genes *TNC* (**C**), *ATP1A2* (**C**, D), and *PTPRZ1* (**D**), together with *DCX* (**C**), which labels DENs, in a human GW23 LGE/MGE sample. Scale bars, 100 μm. **E**) Immunostaining of the cortical tri-IPC markers EGFR and OLIG2 in a human GW14 LGE/MGE sample.

**Supplementary Figure 7.**
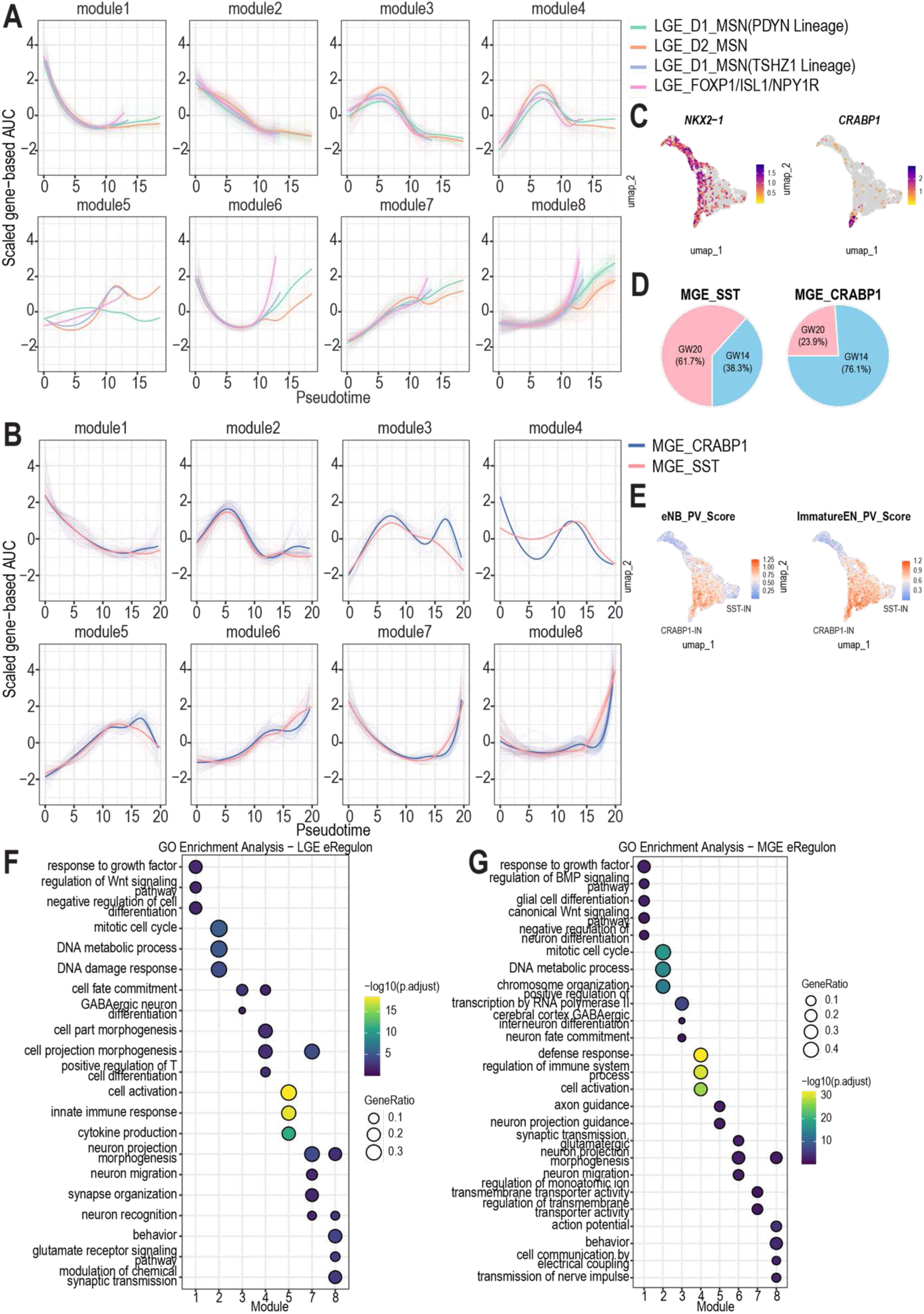
eRegulon-Inferred Developmental Trajectories of LGE and MGE-IN Lineages. **A**, **B**) Standardized gene-based AUC scores of ten eRegulon modules along the LGE (**A**) and MGE-IN (**B**) developmental trajectories. eRegulons are color-coded according to neuronal trajectory, and thick lines indicate trajectory-specific mean AUC values. **C**) UMAP visualization of *NKX2-1* and *CRABP1* expression along MGE trajectories. **D**) Pie charts showing the relative contributions of GW14 and GW20 cells within the MGE_SST_IN and MGE_CRABP1_IN clusters. **E**) UMAP visualization of gene scores for gene sets enriched in early postmitotic neuroblasts (eNBs) of a parvalbumin (PV) fate and immature neurons of a PV fate, as defined in Ma et al., 2021, mapped onto MGE trajectories. **F**, **G**) GO enrichment analysis of target genes associated with each eRegulon module in the LGE (**F**) and MGE-IN (**G**) developmental trajectories. Blank areas denote terms with adjusted P > 0.05. Statistical significance was evaluated by one-sided hypergeometric testing followed by Benjamini–Hochberg adjustment.

**Supplementary Figure 8.**
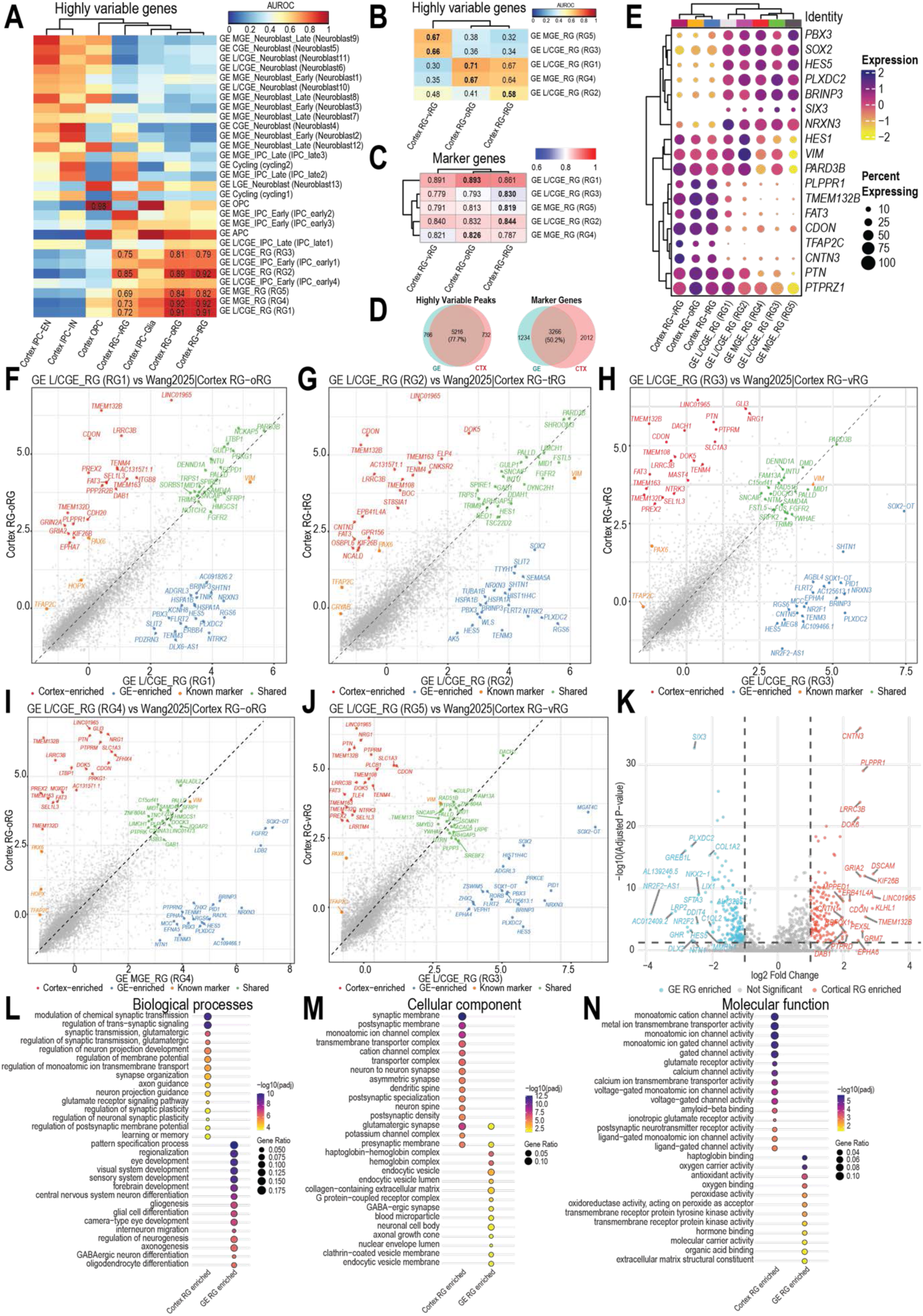
Transcriptomic Comparison of GE RG with Cortial RGs. **A**, **B**) Heatmaps showing MetaNeighbor AUROC-based transcriptomic similarity between cortical and GE progenitor subtypes (**A**) and a zoomed-in comparison of cortical and GE RG types (**B**). Red indicates high similarity, and blue indicates low similarity. **C**) Heatmap showing Spearman correlation scores based on marker gene expression between cortical and GE progenitor subtypes, with a zoomed-in view on RGs. **D**) Venn diagrams showing the overlap between highly variable genes identified by MetaNeighbor and marker genes identified by Seurat differential expression analysis (MAST; log₂FC ≥ 0.25, padj ≤ 0.01). **E**) Clustered dot plot of common and regional-enriched genes between cortical and GE RG. **F**-**J**) Scatter plot showing z-scores of highly variable genes between the most similar pair of cortical and GE RG subtypes. Z-scores were scaled within each subtype. Colors indicate gene categories: red and blue, top 20 most divergent genes in cortical and GE RG, respectively; green, top 20 most similar genes; orange, known cortical RG subtype genes. **K**) Volcano plot showing differential gene expression between cortical and GE RG. Genes with |log₂FC| ≥ 1 and p < 0.05 are highlighted. Red indicates genes upregulated in cortical RG, and blue indicates genes upregulated in GE RG. **L**-**N**) GO enrichment analysis of genes enriched in cortical and GE RGs, showing biological processes (**L**), cellular components (**M**), and molecular functions (**N**).

**Supplementary Figure 9.**
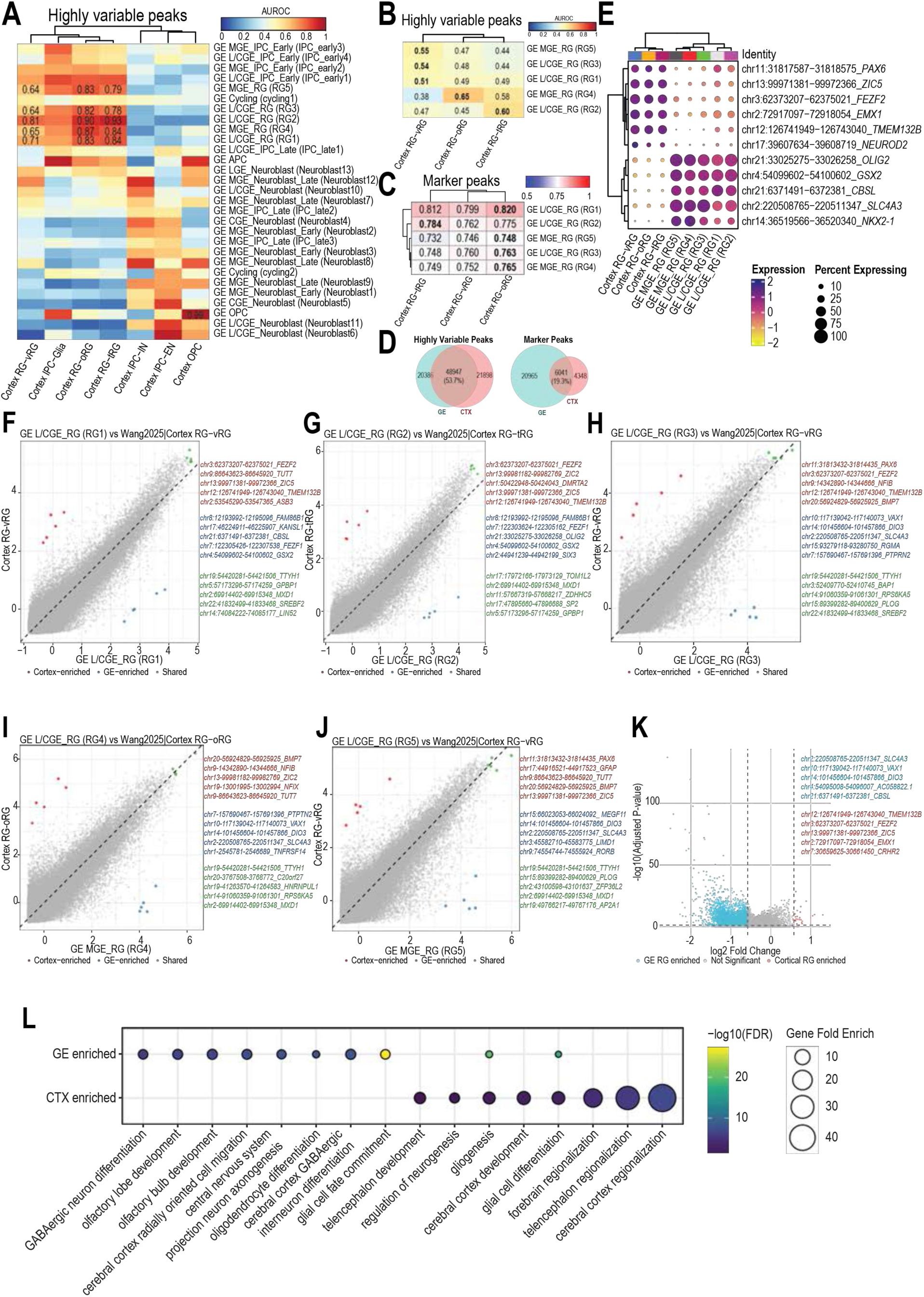
Epigenomic Comparison of GE RG with Cortial RGs. **A**, **B**) Heatmaps showing MetaNeighbor AUROC-based epigenomic similarity between cortical and GE progenitor subtypes (**A**) and a zoomed-in comparison of cortical and GE RG types (**B**). Red indicates high similarity, and blue indicates low similarity. **C**) Heatmap showing Spearman correlation scores based on marker peaks between cortical and GE progenitor subtypes, with a zoomed-in view on RGs. **D**) Venn diagrams showing the overlap between highly variable peaks identified by MetaNeighbor and marker peaks identified by Seurat differential expression analysis (LR; log₂FC ≥ 0.25, padj ≤ 0.01). **E**) Clustered dot plot of regionally enriched peaks between cortical and GE RG. **F**–**J**) Scatter plots showing z-scores of highly variable peaks between the most similar pair of cortical and GE RG subtypes, scaled within each subtype. Top common and divergent peaks are highlighted: red and blue, top 5 peaks specific to cortical and GE RG, respectively; green, top 5 peaks shared between cortical and GE RG. **K**) Volcano plot showing differential chromatin accessibility between cortical and GE RG. Peaks with |log₂FC| ≥ 0.58 and p < 0.05 are highlighted; red indicates peaks upregulated in cortical RG, and blue indicates peaks upregulated in GE RG. **L**) Dot plot showing the most enriched GO terms obtained from peaks enriched in cortical and GE RG.

**Supplemental Figure 10.**
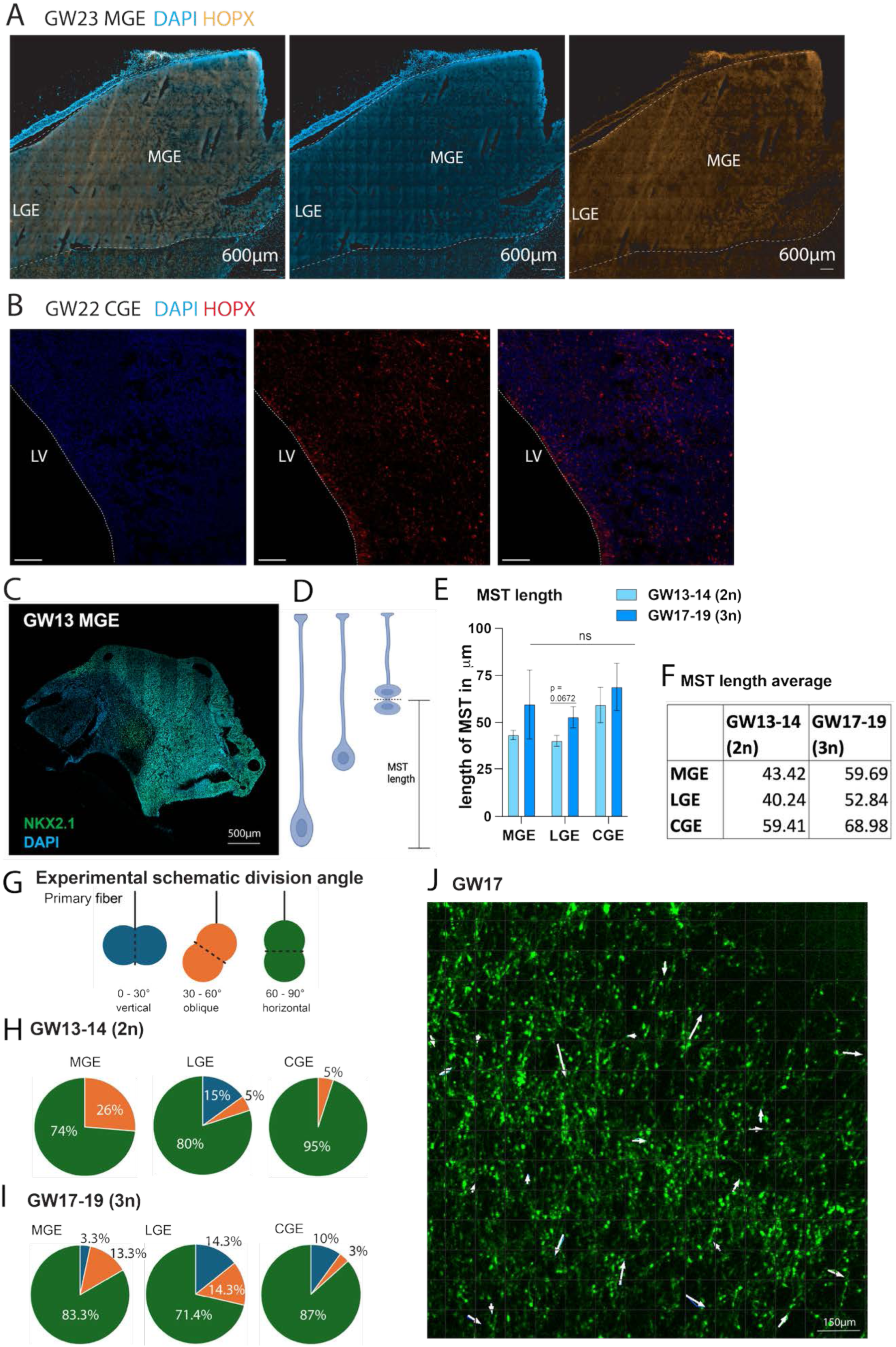
Characterization of division dynamics in the developing human GEs. **A-B)** Immunostaining of HOPX in a GW23 MGE (**A**) and GW22 (**B**) CGE sample. **C)** Immunostaining of NKX2.1 for MGE-identity validation of live imaged organotypic slice at GW13. **D)** Schematic of MST length measurement, with MST length being the translocation of cell body (“jump”) immediately preceding cytokinesis. **E)** MST length at GW13-14 (2n) and GW17-19 (3n) in μm. **F)** Average MST length across GE subregions and ages. Unit is μm. **G)** Division angle analysis of MST-exhibiting cells. Schematic of categorization of division angles with respect to primary fiber: 0-30° (vertical), 30-60° (oblique) and 60-90° (horizontal). **H)** Proportions of division planes in human MGE, LGE and CGE at GW13-14 (2n). **I)** Proportions of division planes in human MGE, LGE and CGE at GW17-19 (3n). **J)** Directionality of MST jumps in GW17 is random. Each white arrows indicates a cell undergoing MST.

**Supplemental Figure 11.**
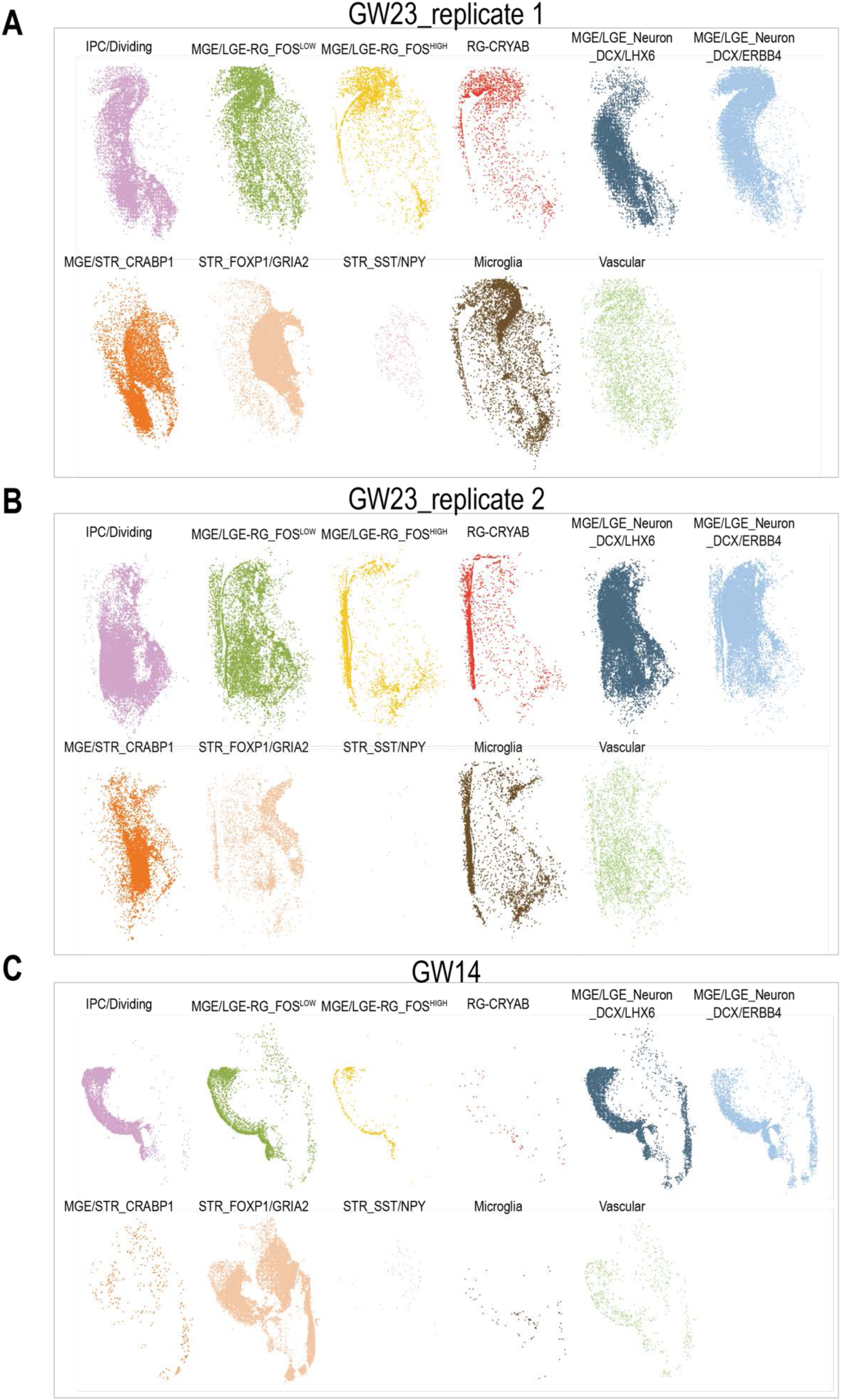
Spatial distribution of cell types in individual MERFISH experiments. **A–C**) Spatial distribution of cell types in GW23 replicate 1 (**A**), GW23 replicate 2 (**B**), and GW14 (**C**).

**Supplemental Figure 12.**
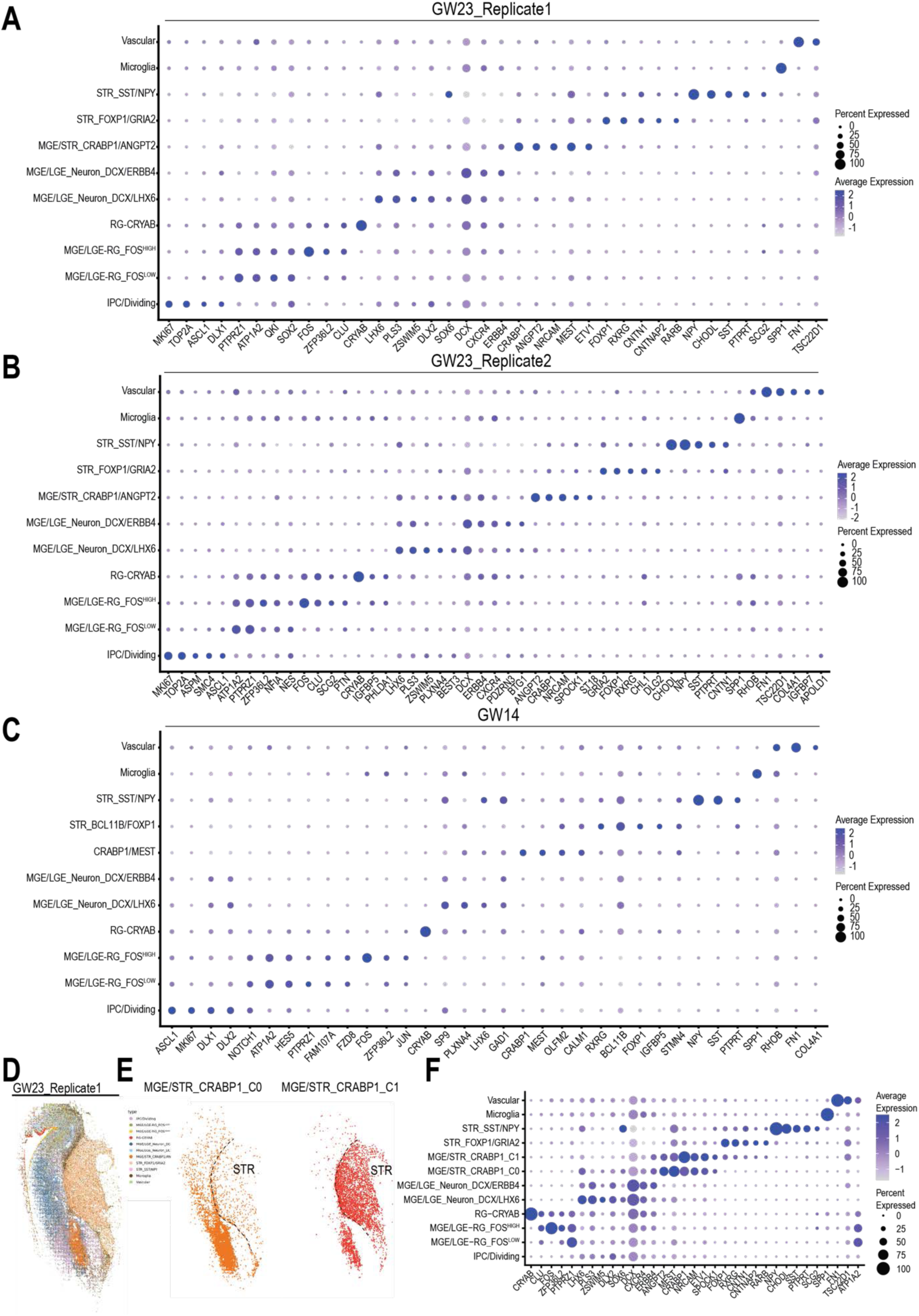
Gene expression patterns and subclustering analysis of the spatial transcriptomic data. **A–C**) Dot plots showing marker gene expression for each cell cluster in GW23 replicate 1 (**A**), GW23 replicate 2 (**B**), and GW14 (**C**). **D**) Spatial transcriptomic map of GW23 replicate 1, with cells color-coded by cell type. **E**) Subclustering of the MGE/STR_CRABP1 cluster. **F**) Dot plots showing marker gene expression for each subcluster identified in the subclustering analysis.

**Supplemental Figure 13.**
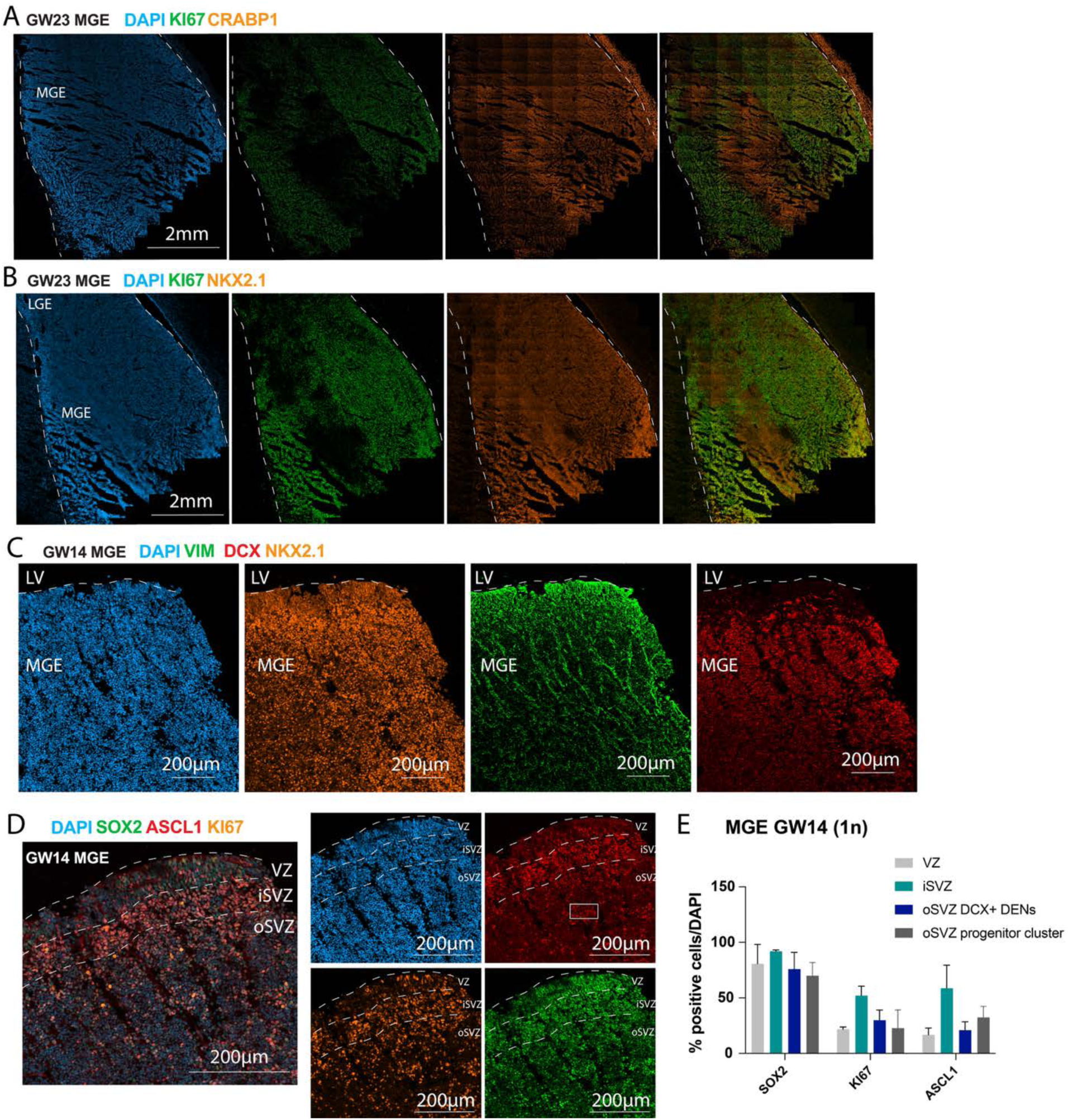
Spatial domains within the human MGE. **A-B)** Immunostaining of KI67+ and CRABP1 (**A**) and NKX2.1 (**B**) reveals that a CRABP1+ cell cluster is located within the human MGE (GW23) and depleted of dividing cells. **C)** Immunostaining DCX+ DENs structures within a fixed GW14 human MGE sample from which a consecutive section was used for spatial transcriptomics. **D)** Immunostaining shows enrichment of ASCL1+ IPCs in the iSVZ. White box indicates clusters of ASCL1+ IPC within a column of RG fibers supporting multilevel DEN separation as seen in EM. **E**) Quantification of progenitor markers within MGE subdomains based on immunostaining.

**Supplemental Figure 14.**
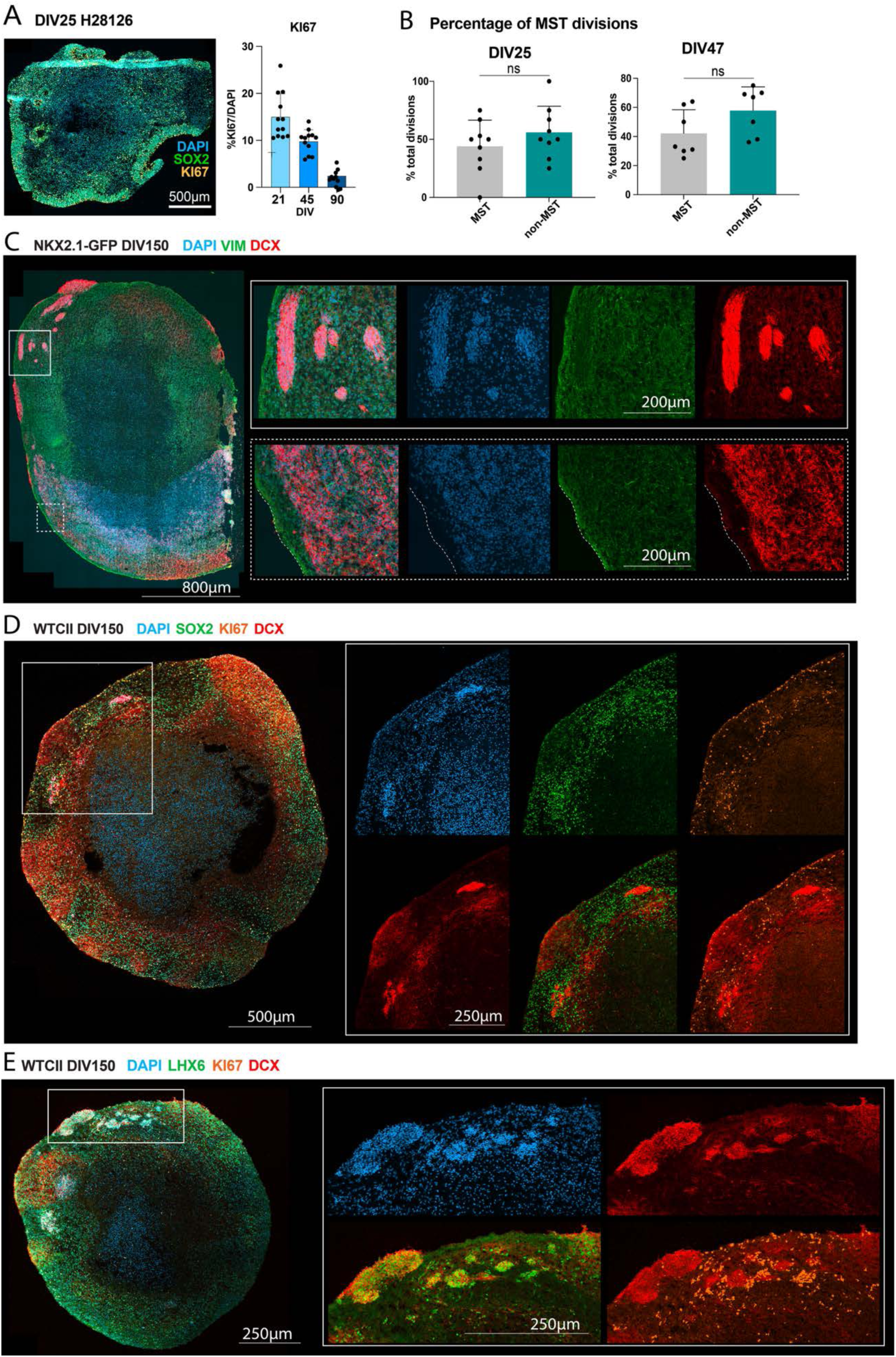
DENs in MGE-enriched regions of GE-organoids. **A)** Immunostaining and quantifications show reduction in KI67+ cells as organoids mature. **B)** Quantifications of division proportions in GE organoids at DIV25 and DIV47 using live imaging and imaris. **C-E)** Immunostaining of MGE-enriched organoids at DIV150 shows mature DENs (**C),** sustained SOX2 and KI67 expression surrounding DENs (**D**), and DEN areas express LHX6 indicating MGE identify (**E**).

## STAR Methods

### KEY RESOURCES TABLE

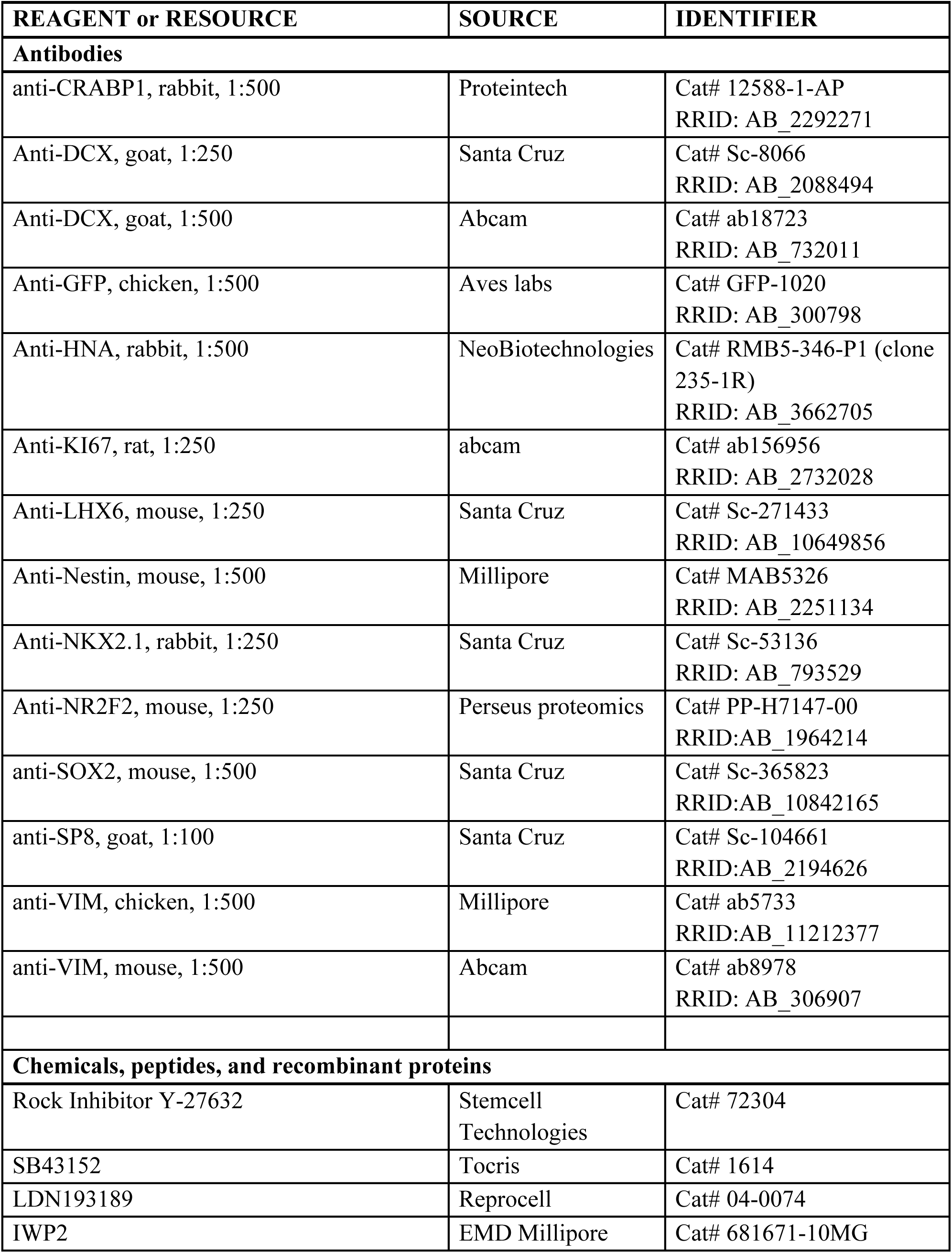
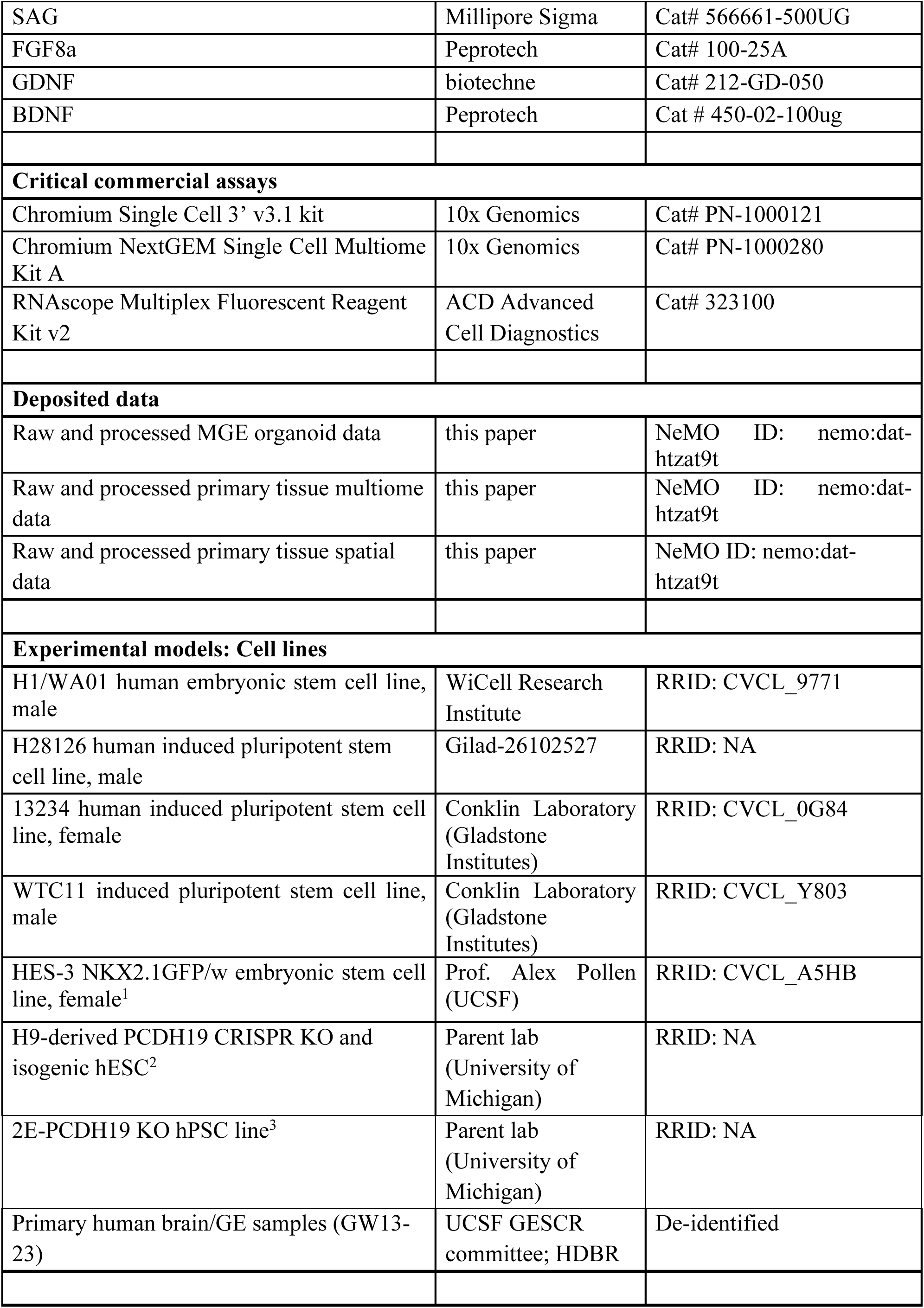
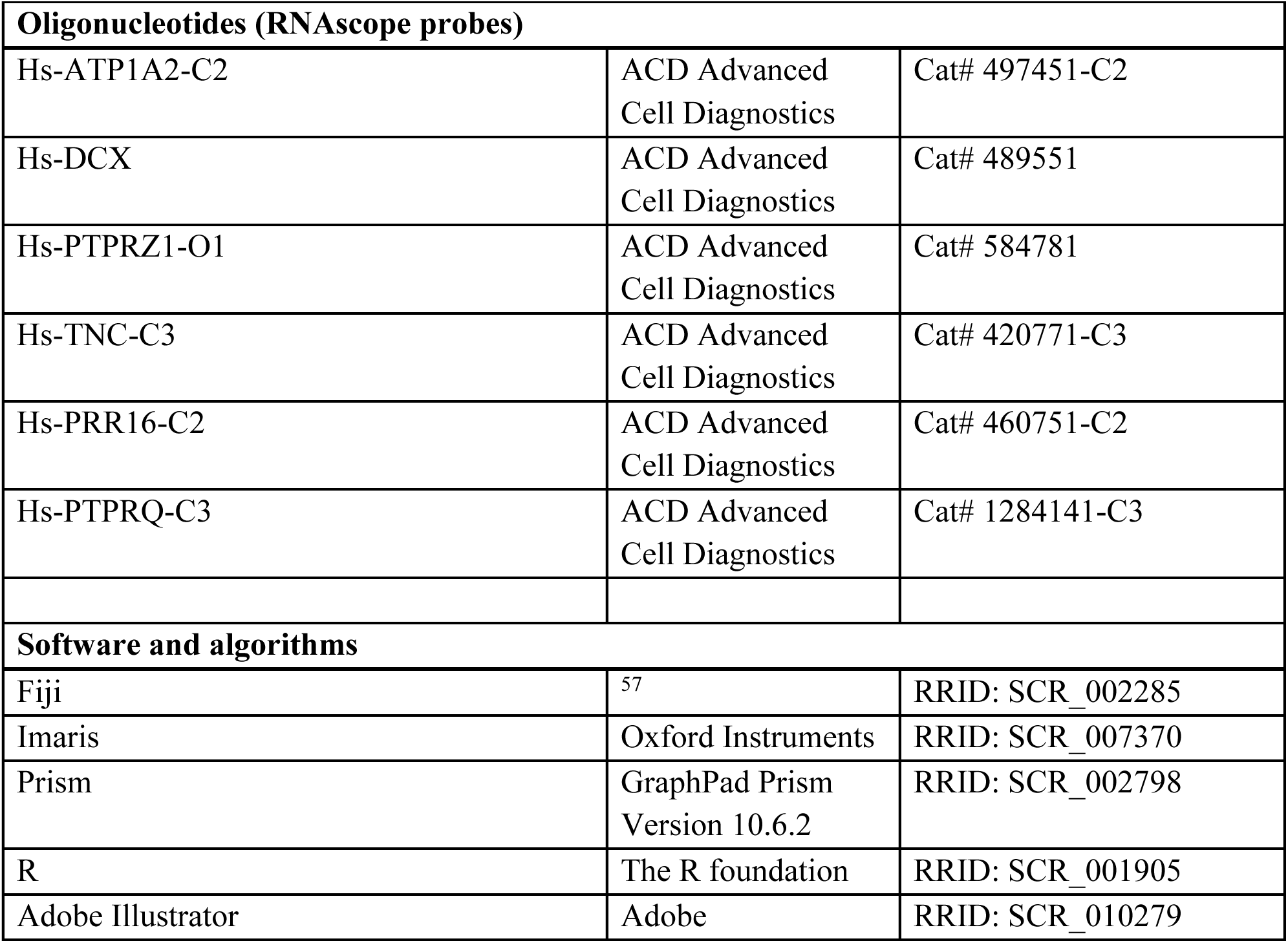

## Resource availability

### Lead contact

Further information and requests for resources and reagents should be directed to and will be fulfilled by the lead contact, Arnold Kriegstein.

### Materials availability

This study did not generate new unique reagents.

## Experimental models and study participant details

### Pluripotent Stem Cell Lines

Previously described unmodified human PSC lines H1/WA01, WTCII, 13234, H28126, in addition to the transgenic NKX2.1-GFP+ reporter cells line^1^ and female H9-derived PCDH19 CRISPR KO^2^, male 2E-PCDH19 KO hPSC line^3^ and isogenic control lines hESC were used. Before use and every 10 passages, cells are tested for karyotypic abnormalities and validated for pluripotency markers Sox2, Nanog and Oct4. All cell lines tested negative for mycoplasma.

### Human pluripotent stem cell culture

Human pluripotent stem cells (hPSCs), frozen in mFreSR (Stem Cell Technologies; #5855), were thawed and cultured on growth-factor reduced Matrigel (BD Biosciences; #354230)-coated six well plates in Stem Flex Pro Medium (Gibco) with 10 μM Rock inhibitor Y-27632. Medium was changed every other day without Rock inhibitor until about 70% confluency was reached. For maintenance passaging, PBS-EDTA (Lonza) was added to cells for 1 min, aspirated, and cells were incubated at 37°C for 3 mins. Cells were resuspended in Stem Flex Pro Medium and plated into new growth-factor reduced Matrigel-covered plates.

### MGE organoid differentiation protocol

hPSCs were differentiated into cortical organoids based on published protocols for directed forebrain differentiation^4,5^. All medium and small molecules were based on the previously published protocol^4,5^, just the aggregation and culture conditions were changed. On Day 0, PSCs were dissociated into single cells using accutase and plated into 96-well ultra-low attachment v-bottom plates at about 15,000 cells per well. Cells were aggregated in KSR medium composed of DMEM (Gibco; #11995-065) containing 15% v/v Knockout Serum Replacement (Lifetech; #10828-028), 2mM L-glutamine (Gibco, #A2916801), 10μM ß-mercaptoethanol (Sigma) and 1x Primocin (InvivoGen; #ant-pm-1). Fresh TGF-B inhibitor SB431542 (10μM; Tocris; #1614), LDN193189 (100nM; Reprocell; #04-0074), IWP2 (5μM; EMD Millipore; #681671-10MG) and Smoothened agonist SAG (0.1μM; Millipore Sigma; #566661-500UG) were added at every medium change until DIV8. Rock inhibitor Y-27632 (20μM; Tocris; #1254) was added only on day 0. From DIV8-14 only LDN193189 (100nM) and Smoothened agonist SAG (0.1μM) were added. Medium was changed every other day (3 times a week). Once stable cell aggregates/spheres had formed (between DIV7-10), organoids were transferred to low adhesion 6-well plates and moved to an orbital shaker rotating at 90 rpm. On DIV14, the culture medium was changed to N2AA medium consisting of DMEM/F-12 (Gibco; #10565-042) containing 0.5x/1:200 N2 supplement (Gibco; #17502-048), Ascorbic Acid (200μM; Sigma-Aldrich; #A0278-25G) and 1x Primocin. From DIV14-21, SAG (0.1μM) and FGF8a (100 ng/ml; Peprotech; #100-25A) were added to the N2AA medium. From DIV21-45, glia cell line-derived neurotrophic factor (GDNF; 10 ng/ml; biotechne; #212-GD-050) and brain-derived neurotrophic factor (BDNF; 10 ng/ml; Peprotech; #450-02-100ug) were added. At DIV45, medium was switched to B27GB medium consisting of DMEM/F12 with 0.5x/1:100 B27 supplement (Gibco; #17504-044), BDNF (10 ng/ml), GDNF (10 ng/ml) and 1x Primocin.

For PCDH19-KO organoids, female H9-derived PCDH19 CRISPR KO^6^, male 2E-PCDH19 KO hPSC lines^3^ and isogenic control lines were differentiated into MGE regional specific brain organoids as described in our preprint^6^.

### Primary human brain tissue

Primary human brain tissue of the cortex and ganglionic eminences was obtained and processed as approved by UCSF Gamete, Embryo and Stem Cell Research Committee (GESCR). All samples were de-identified including information on sex. Tissue was collected with informed patient consent for research and in strict observance of legal and institutional ethical regulations. Samples were obtained from either the Human Developmental Biology Resource (HDBR) (hdbr.org) (age was determined using crown-rump length) or the Zuckerberg San Francisco General Hospital (ZSFGH).

### Primary tissue organotypic slice cultures

Primary human GE tissue obtained from ZSFGH was processed within 12hrs postmortem in artificial cerebrospinal fluid (ACSF) containing 125 mm NaCl, 2.5 mm KCl, 1 mm MgCl2, 2 mm CaCl_2_, 1.25 mm NaH_2_PO_4_, 25 mm NaHCO3, 25 mm D-(+)-glucose. ACSF was bubbled with a 95% O2 - 5% CO2 gas mixture before use. Human GE tissue obtained from HDBR was shipped in Hibernate-E medium (Gibco; #A1247601) with B27 and PS at 4°C. Tissue was received at UCSF 2-3 days postmortem and processed immediately.

For organotypic slices, tissue was embedded in 3.5% w/v low melt agarose (Fisher scientific; BP165-25) and sectioned into 300μm slices using a vibratome (Leica). Floating slices were incubated in ACSF with 1:2000 Ad-CMV-GFP (vectorbiolabs; #1060) for 30 min. at 37°C. Slices were transferred to Millicell inserts in a 6 well plate and cultured at the air liquid interface with slice culture medium (66% v/v Basal Medium Eagle, 25% v/v Hank’s Balanced Salt solution, 5% v/v FBS, 0.67% w/v glucose, 1x Penicillin-Streptomycin-Glutamine, 1x N2 supplement).

## Methods details

### Organoid time-lapse imaging

Live imaging was conducted using 4 cell lines (WTCII, H1, H28126 and 13234). Organoids were embedded in 5% w/v low melt agarose (Fisher scientific; BP165-25) in PBS and sectioned into 500 μm slices using a vibratome (Leica). Slices were incubated floating in cell culture medium according to age of organoids with 1:1000 Ad-CMV-GFP (vectorbiolabs; #1060) at 37°C for 3-5 days. One day after adding the virus, the same volume used for infection of slice culture medium (66% v/v Basal Medium Eagle, 25% v/v Hank’s Balanced Salt solution, 5% v/v FBS, 0.67% w/v glucose, 1x Penicillin-Streptomycin-Glutamine, 1x N2 supplement) without virus was added to organoid slices without removing any medium. Once solid GFP expression was observed (usually 3 days post infection), a full medium change was conducted, and organoid slices were transferred to Millicell inserts in a 6 well plate to be cultured at the air liquid interface with slice culture medium. Organoid slices were acclimated to slice culture medium for at least 3 days before live imaging was started. Millicell inserts with slices were transferred to a glass-bottom plate and imaged for 72 hrs using an inverted Leica TCS SP5 or a Zeiss LSM900 with an incubation chamber attached that allowed continuous gas (5% CO2, 5% O2, balance N2) and temperature control (37°C). GFP+ cells were imaged using a 10x air objective at 20-30 min intervals. The focus was checked twice daily, and z-steps were adjusted if needed to keep cells in focus.

### Primary tissue time-lapse imaging

Once stable GFP expression was observed (24-48 hours after viral labelling), live imaging was started. Slice inserts with slices were transferred to glass-bottom plates with fresh medium and imaged for 72 hrs using an inverted Leica TCS SP5 with an incubation chamber attached that allowed continuous gas (5% CO2, 5% O2, balance N2) and temperature control (37°C). GFP+ cells were imaged using a 10x air objective at 20-30 min intervals. The focus was checked twice daily, and z-steps were adjusted if needed to keep cells in focus.

### Transplantation of GE organoids into mouse cortex

Organoids at DIV130 were dissociated using the Papain Dissociation System (Worthington Biochemical). Organoids were incubated in papain solution at 37°C for 30 mins on a shaker. The reaction was quenched with an equal volume of culture medium, and organoids were gently triturated using pipettes to obtain a single-cell suspension. Cells were washed twice with L-15 medium (gibco; #11415-004) supplemented with DNase I (60U/ml), filtered through a 40-μm strainer, and resuspended at 50,000 cells/μl in L-15 medium with DNase I. The cell suspension was loaded into mineral oil–prefilled bevelled glass micropipettes (70–90 μm, Wiretrol 5 μl, Drummond Scientific) mounted on a microinjector. Postnatal day 5-6 NSG mice (JAX; #005557) were anesthetized by hypothermia (∼4 min) and placed in a clay head mold for skull stabilization. Using a stereotactic injector, beveled glass micropipettes were positioned vertically above the target site. Injections were carried out bilaterally, with entry points aligned perpendicular to the skin surface at eye-level coordinates (x = 1, y = 4). A total of 50 nl of cell suspension was dispensed at successive depths of 0.2, 0.4, 0.8, and 1 mm below the skin surface. The pipette was left in place for 1 minute following the final deposition before being gently withdrawn. Pups were placed on a pre-warmed heat pad for recovery before being returned to their mothers. At 52 days post-transplantation, mice were euthanized and transcardially perfused. Brains were obtained, fixed, cryoprotected, and sectioned at 50 μm thickness using a cryostat, followed by phenotypic analysis through immunostaining.

### Single-cell RNA sequencing (ScRNAseq) of organoids

For scRNAseq, organoids produced from the cell lines H28126, H1 and WTCII were sequenced at DIV51 and DIV164. For the cell line 13234, we only sequenced DIV51. MGE organoids were dissociated using the papain dissociation system (Worthington; #LK003150) for 30-40 min at centrifuged at 300g for 5 min. Organoids were washed twice in PBS + 0.1% BSA and passed through a FACS filter. Single-cell capture was performed using the 10xGenomics Chromium Single Cell 3’ v2 kit per manufacturer’s recommendations. Single-cell libraries were sequenced on a NovaSeq S2 flow cell. All cell lines of the same age were mixed and ran in two lines. At this point in the analysis, the cell lines of origin had not yet been demultiplexed, and all data were treated as a pooled population.

### Multiome sequencing of primary tissue

For nuclei isolation, 25–35 mg of fresh-frozen tissue was processed for nuclei extraction following a published protocol^7^. Counted nuclei were subjected to transposition using the Chromium Next GEM Single Cell Multiome platform and subsequently loaded onto a 10x Genomics chip for GEM generation according to the manufacturer’s instructions. Libraries from individual samples were pooled and sequenced on a NovaSeq X system, targeting approximately 25,000 read pairs per nucleus for RNA and 50,000 read pairs per nucleus for ATAC.

### Nanostring GeoMx sample preparation

Nanostring GeoMx was run using one biological replicate (GW23 MGE) according to the manufacturer’s recommendation (MAN-10150-02, p.50, then p.35-p.44). In short, 10mm sections were mounted onto Superfrost™ Plus Microscope Slides and frozen at -80°C. Slides were processed according to the “Fresh frozen sample preparation for RNA assays” protocol (p.50, MAN-10150-02) with a 3 hrs baking step. Target retrieval was performed according to the “Perform target retrieval” protocol (p.35, MAN-10150-02) with target antigen retrieval for 20 min at 100°C. During the “Expose RNA targets” step, two proteinase K concentrations were used, 1mg/ml and 0.1mg/ml. No post fixation was performed. *In situ* hybridization and stringent washing were performed according to manufacturer’s protocol without changes. For morphology markers, Syto13 and the conjugated antibodies DCX-594 (1:200, santa cruz biotechnology, #sc-271390 AF594), VIM-PE (1:400, #sc-373717 (Vim E-5) PE), and KI67-647 (1:100, abcam, #ab196907) were incubated for 1 hr at RT. The sample was run on the Nanostring GeoMx platform according to MAN-10152-02 and the post-run PCR and sample preparation for sequencing was performed using MAN-10133-05. Samples were sequenced on the NovaSeq PE150 platform by novogene.

### Vizgen Merscope gene panel design

A 433-gene custom gene panel was created by integrating single cell and/or single nucleus RNA sequencing data of MGE, LGE and cortical samples from available datasets^8–10^. Top 10 cell type markers were obtained by integrating single cell RNA sequencing data of MGE and LGE samples. Normalization, integration, variable features detection and top marker identification were performed using the standard settings of the Seurat v.5 package (https://satijalab.org/seurat/articles/integration_introduction). CellChat analysis was performed to identify commonly expressed receptor-ligand signaling pathways. Differentially expressed genes between VZ and oSVZ were obtained from the GeoMx pilot dataset of one GW23 MGE sample. Candidate genes from all analyses were compiled, duplicates removed and any genes with high fpkm expression were removed according to manufacturer’s recommendation (https://vizgen.com/gene-panel/), leading to a final count of 433 genes being included in the gene panel (**Table 1**).

**Table 1.**
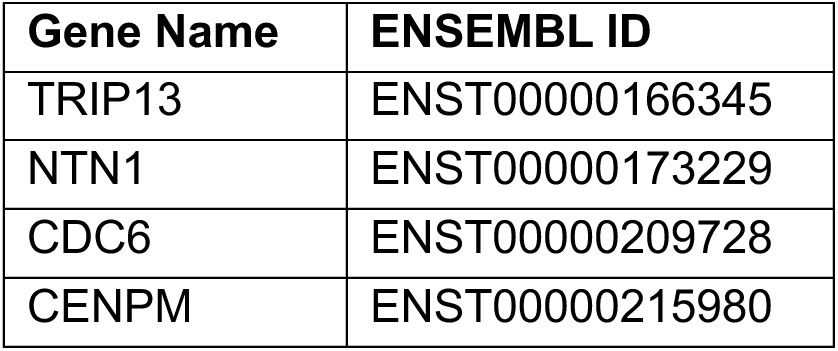
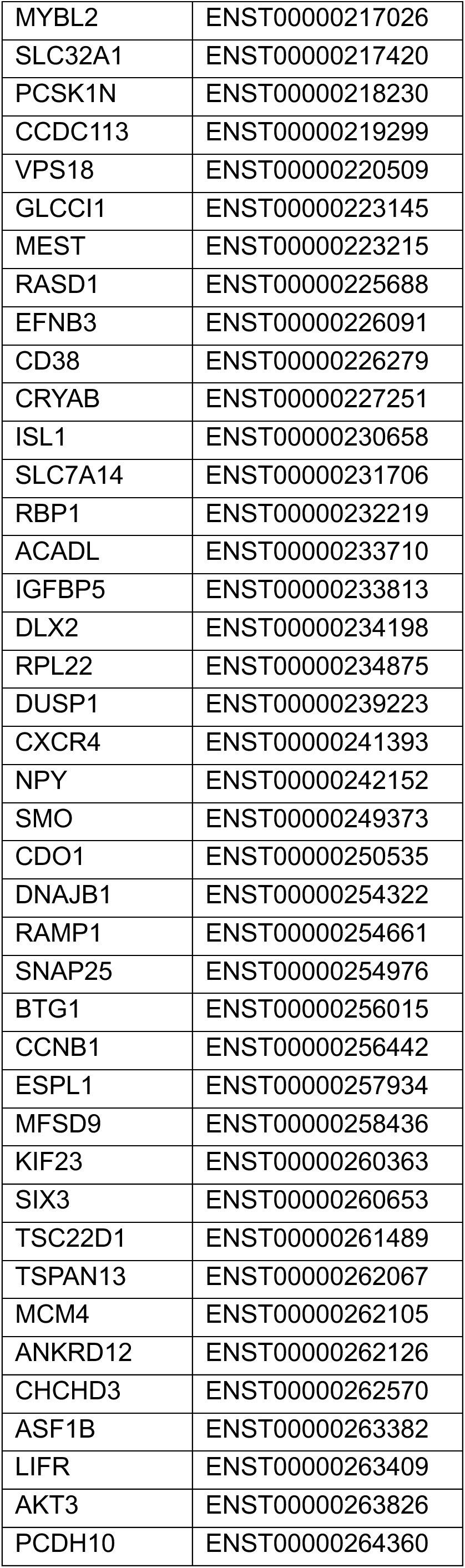
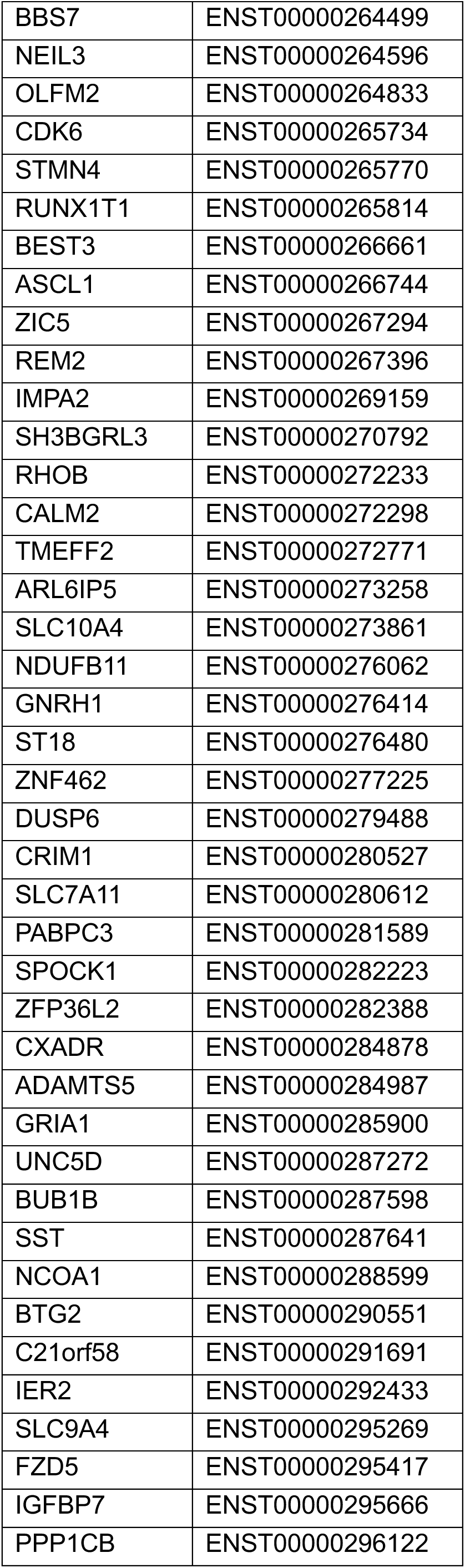
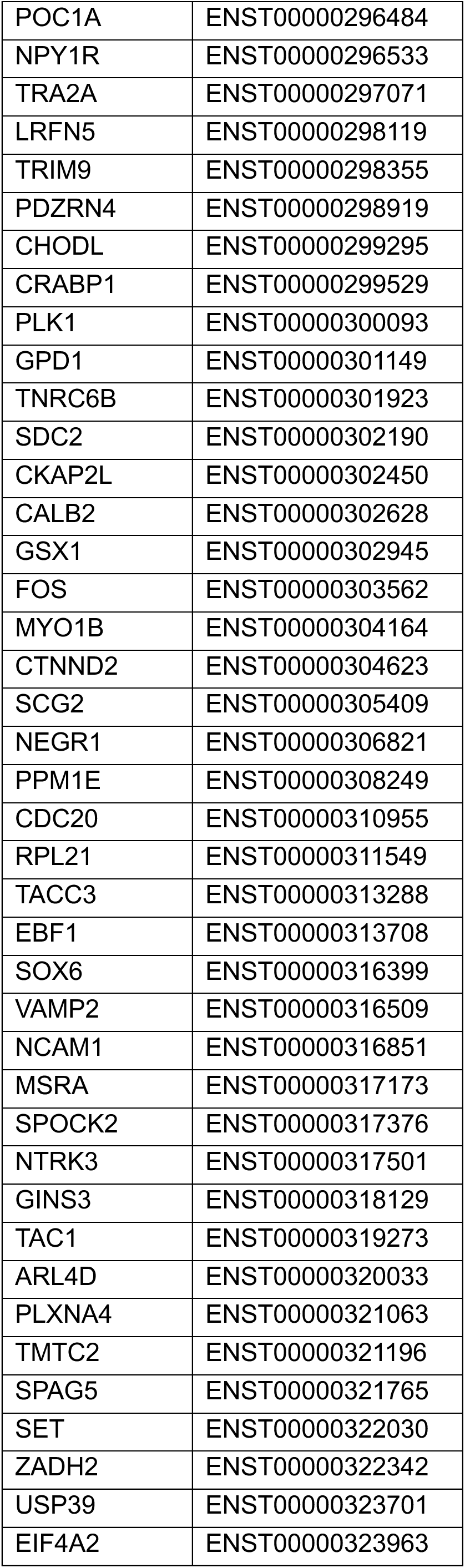
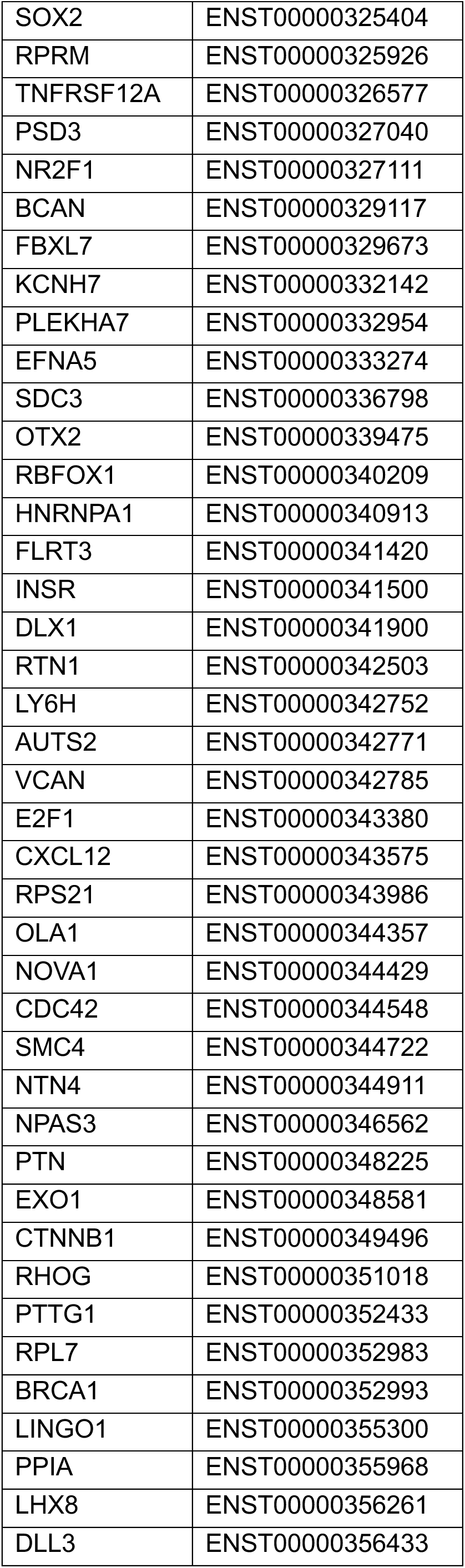
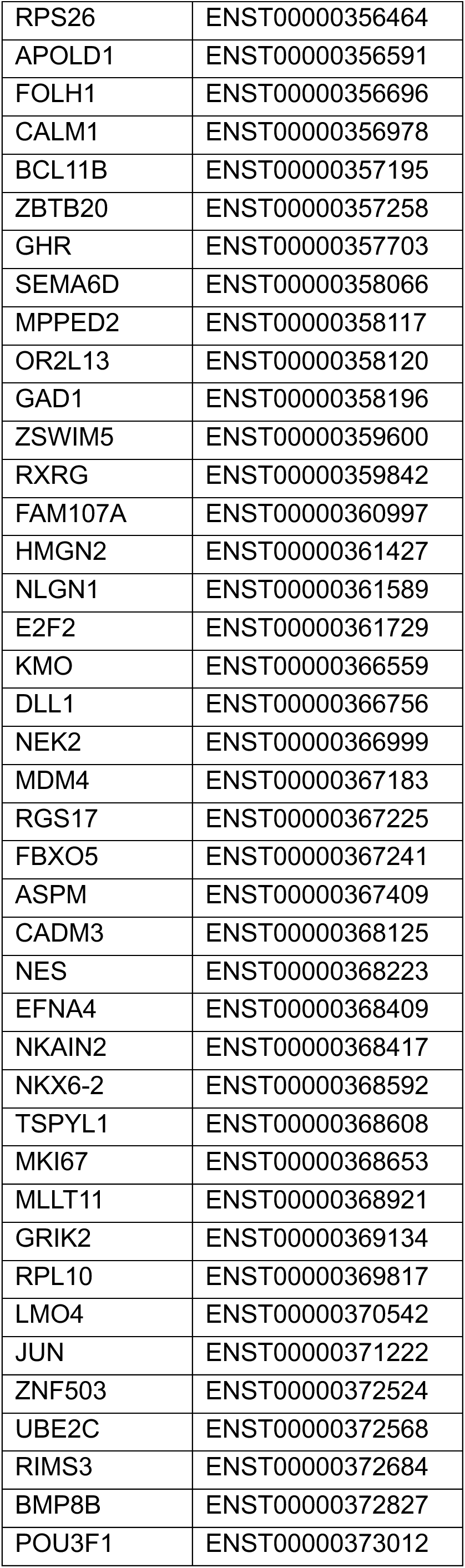
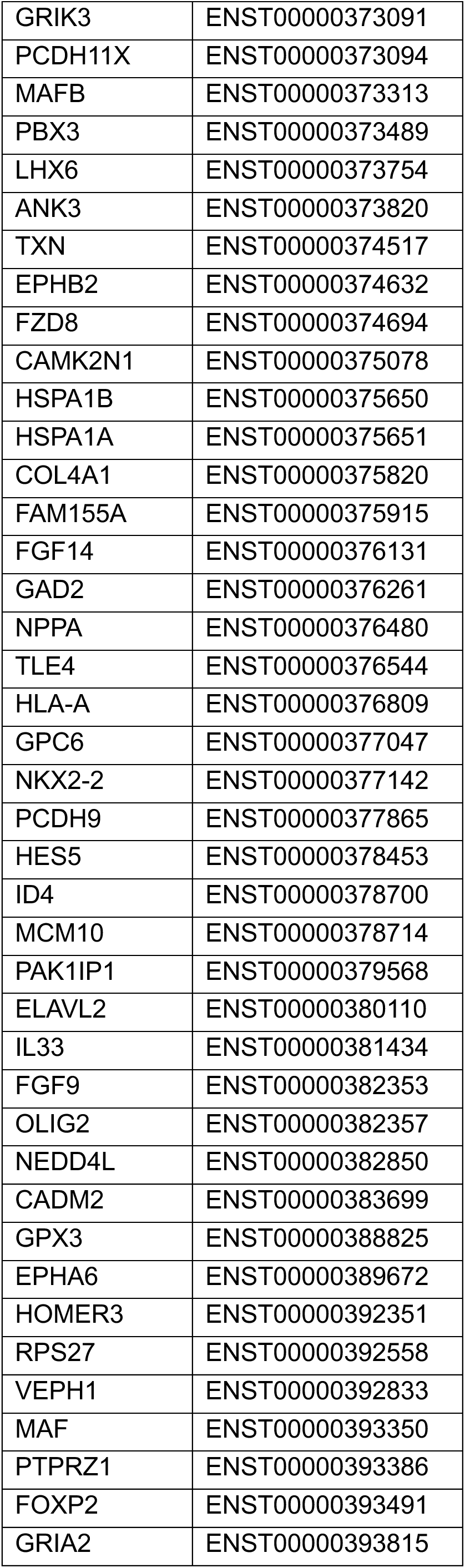
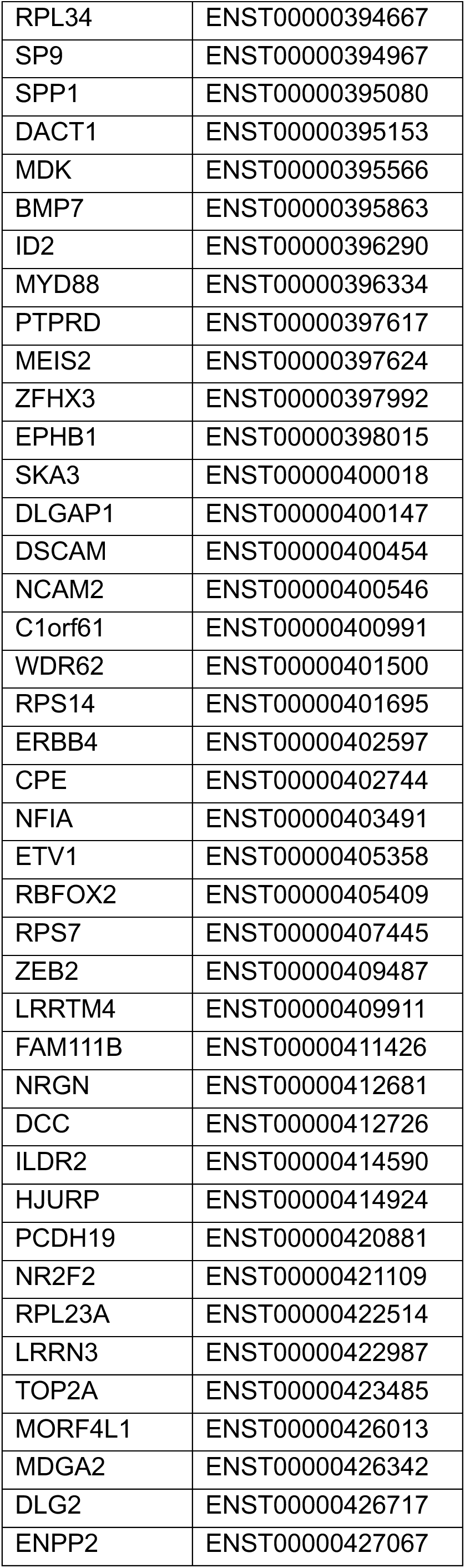
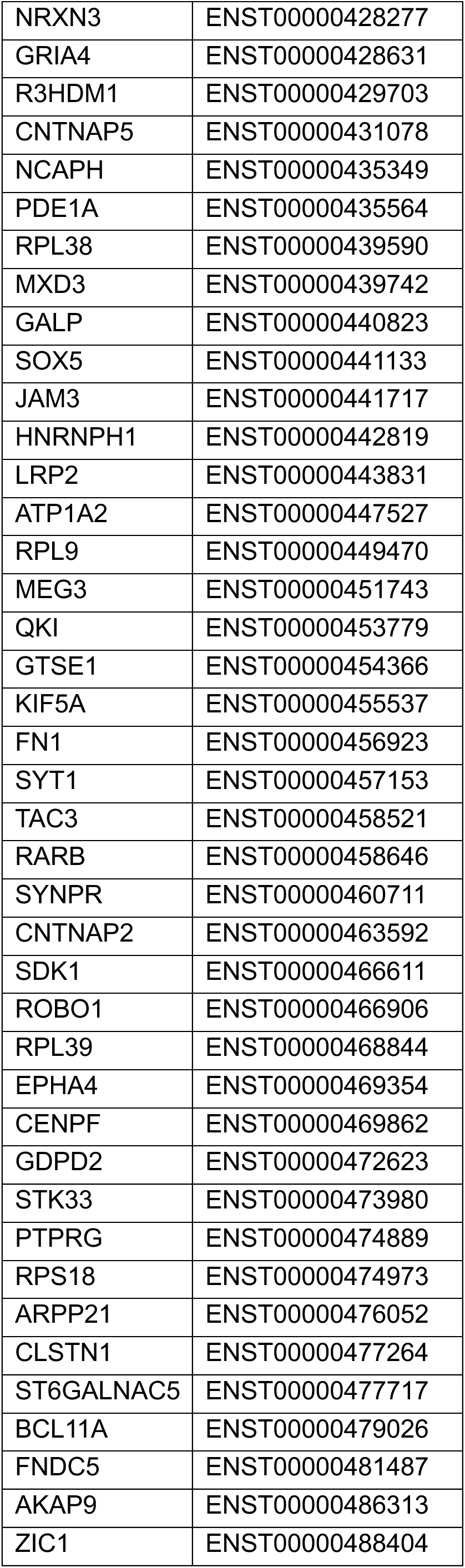
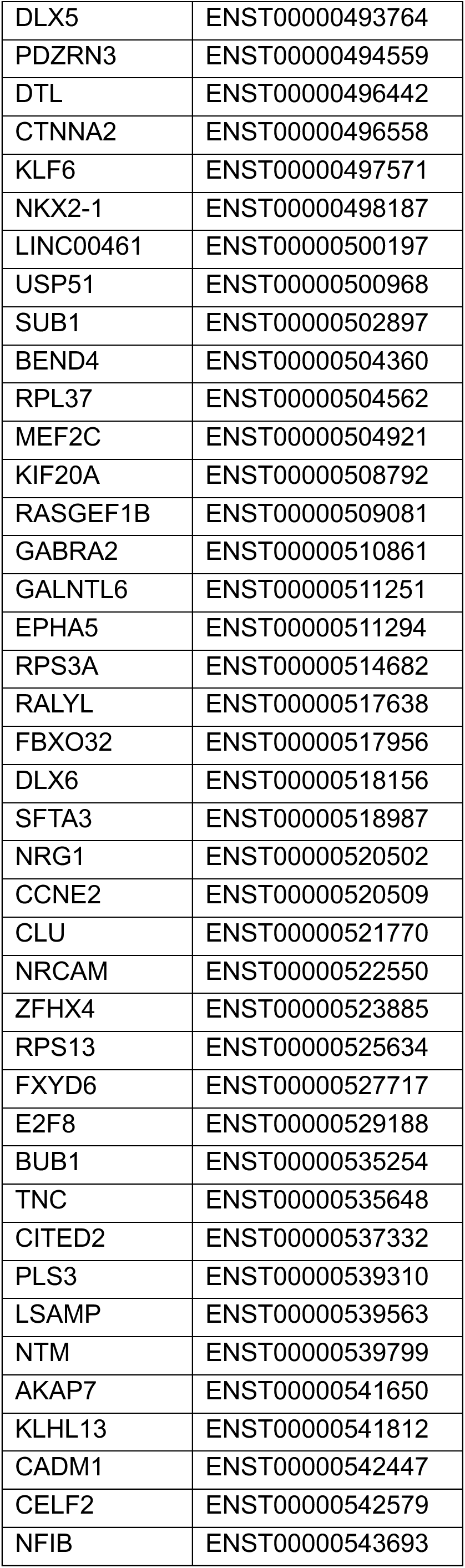
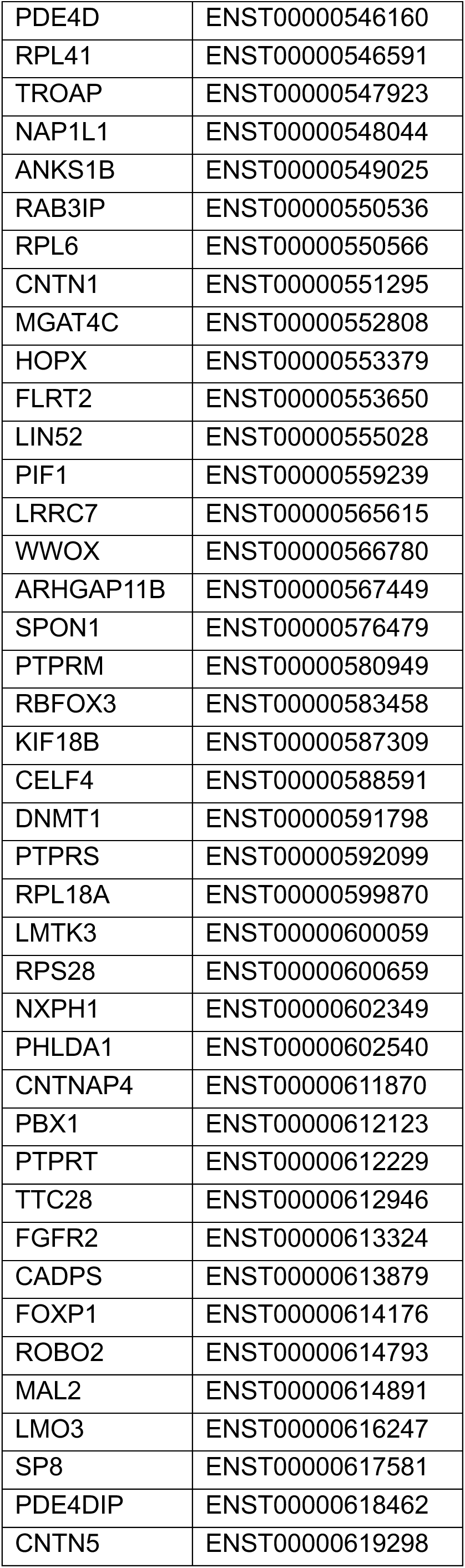
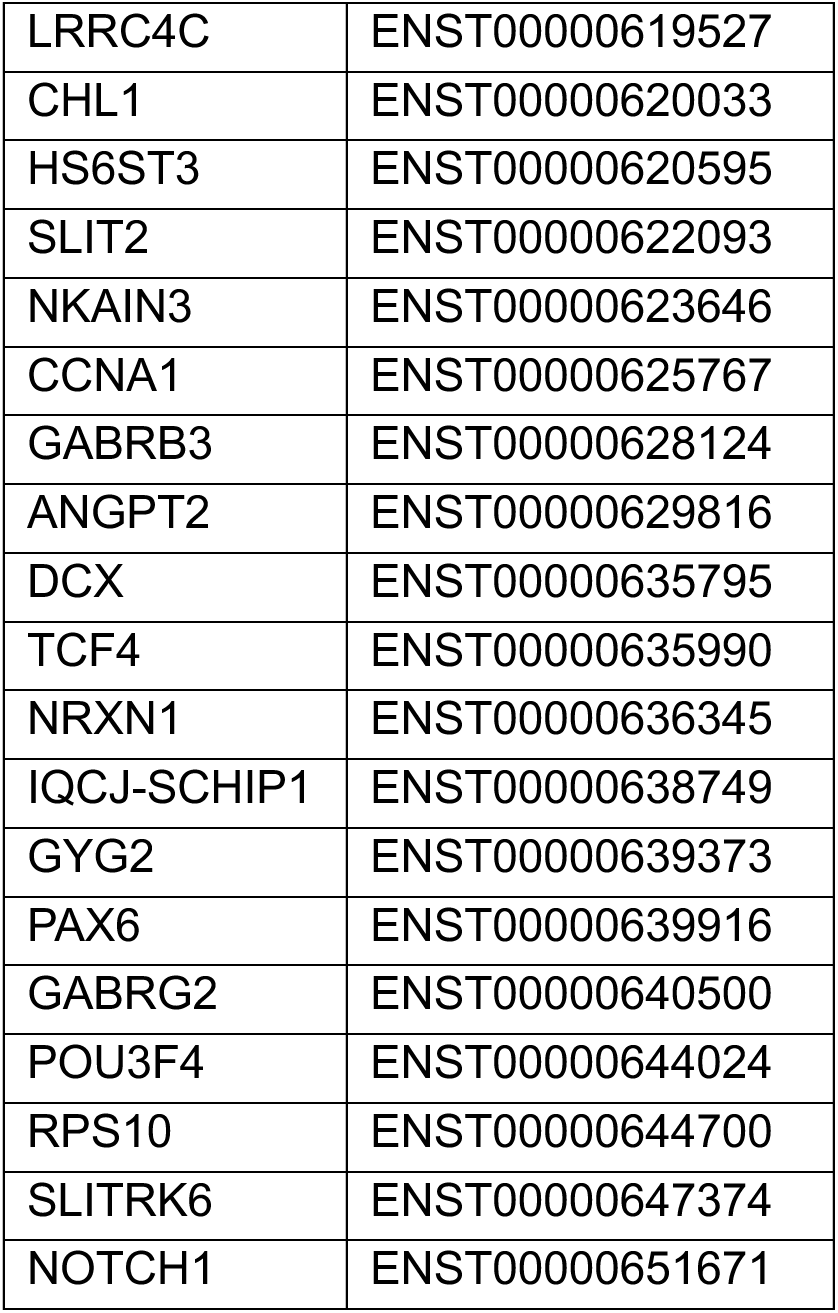
Merscope gene panel of 433 genes.

### Vizgen sample preparation

One fresh frozen GW23 coronal section (MGE and LGE) was prepared as 10mm sections on MERSCOPE slides according to manufacturer’s recommendation (p.24, 91600002 Rev C) including the optional fixation step after sectioning. Then the protocol for cell boundary and protein staining was performed (p.32-40, 91600002 Rev C). For cell boundary staining only the secondary cell boundary was added (primary is incompatible with rabbit antibodies) in addition to goat-DCX (1:250, santa cruz biotechnology, #sc-8066), mouse-Vim (1:500, santa cruz biotechnology, #sc-373717) and rabbit-CD31 (1:500, Invitrogen, #MA5-29475). For secondary staining, Anti-Mouse Aux 4 (PN 20300101), Anti-Rabbit Aux 5 (PN 20300102) and Anti-Goat Aux 6 (PN 20300103) were used. The fixed GW14 sample was processed as the GW23 sample but without the fixation step after sectioning.

### Fluorescent in situ hybridization (RNAscope)

Primary GE tissue and MGE organoids were sectioned into 10mm sections using a vibratome. RNAscope was performed using the RNAscope Multiplex Fluorescent Reagent Kit v2 (ACD Advanced Cell Diagnostics) according to manufacturer’s recommendation. For pretreatment, slices were post-fixed for 10 min. in chilled 4%PFA, washed 2x with MilliQ-H20, antigen retrieval was performed for 5 min., slices were washed 2x with MilliQ-H20 and dipped in 100% ethanol. Protease III was used for 30 min. at 42°C if no co-immunostaining was performed, while Proteinase Plus was used for 10 min. if co-immunostaining was performed. Slides were washed 2x with MilliQ-H20 and probe (see **Key resource table**) hybridization (see Key resource table) was performed. As secondary fluorescent probes, Fluorophores Opal-520 (1:500), Opal-570 (1:750) and Opal-690 (1:750) (all Akoya Biosciences) were used.

### Immunohistochemistry

Intact primary GE samples were fixed in 4% PFA overnight at 4°C, while organoids and organotypic slice cultures were fixed in 4% PFA for 45 mins. Fixed samples were washed with 1x PBS and incubated in 30% sucrose in 1x PBS overnight at 4°C. Samples were embedded in cryomolds in 3:2 O.C.T. (Tissue-tek):30% sucrose in 1x PBS, frozen on dry ice and stored at -80°C. Frozen tissue was sectioned into 10μm sections onto glass slides and stored at -80°C. Antigen-retrieval was performed with 1x boiling citric acid-based antigen unmasking solution (Vector Laboratories) for 20 min. If primary intact human GE was obtained as fresh frozen tissue, it was fresh frozen as slabs and fixed in 4% PFA for 10 mins after sectioning and before antigen retrieval (no sucrose dehydration and OCT embedding). Slides were washed once with MilliQ-H2O and twice with PBS and blocked in Donkey blocking buffer (DBB) containing 5% donkey serum, 2% gelatin and 0.1% Triton X-100 in PBS for 30 mins at RT. Primary antibodies (see **Key resources table)** were incubated in DBB overnight at 4°C, washed three times with 0.1% Triton X-100 in PBS and incubated with AlexaFluor secondary antibody (ThermoFisher and Jackson Labs) and DAPI in DBB for 2 hrs at RT. Images were acquired using a confocal microscope (Leica TCS SP5 or Zeiss LSM900). For PCDH19 KO MGE organoids, staining was based on protocols described in our preprint^11^. For mouse transplant immunostainings, whole mouse brains were dehydrated in sucrose for 1-2 days and sectioned into 50um slices using a vibratome (Leica). Antigen retrieval was performed using citric acid for 30 min at 80°C. Slices were blocked in DBB for 30 mins and primary antibodies were incubated for 72 hrs at 4°C. All other steps were performed as above.

### Electron microscopy

For transmission electron microscopy (TEM), 200 micron sections were prepared using a vibratome and postfixed with 2% osmium tetroxide solution. Dehydration was performed in crescent ethanol concentrations. Sections were stained with 2% uranyl acetate, and finally embedded in araldite resin (Durcupan ACM Fluka, Sigma), and polymerized at 69 °C for 72 hours. 1.5 µm serial semithin sections were obtained and stained with toulidine blue.

## Bioinformatics Analysis

### Multiome Bioinformatic analysis

The Cell Ranger-ARC count pipeline (10x Genomics) was used for cell calling, read alignment, and quality control with the GRCh38 human reference genome (GENCODE v32/Ensembl98). Our filtering thresholds were GW14_LGE (1,000 < UMI_RNA < 50,000, 1,800 < UMI_ATAC < 500,000, nucleosome_signal < 2, TSS enrichment > 1), GW14_MGE (1,000 < UMI_RNA < 80,000, 1,800 < UMI_ATAC < 500,000, nucleosome_signal < 2, TSS enrichment > 1), GW14_CGE (1,000 < UMI_RNA < 80,000, 1,800 < UMI_ATAC < 500,000, nucleosome_signal < 2, TSS enrichment > 1), GW20_LGE (1,000 < UMI_RNA < 80,000, 1,800 < UMI_ATAC < 500,000, nucleosome_signal < 2, TSS enrichment > 1), GW20_MGE (1,000 < UMI_RNA < 80,000, 1,800 < UMI_ATAC < 200,000, nucleosome_signal < 2, TSS enrichment > 1), GW20_CGE (1,000 < UMI_RNA < 60,000, 1,800 < UMI_ATAC < 400,000, nucleosome_signal < 2, TSS enrichment > 1). A total of 20,319 high-quality nuclei were retained for downstream analysis.

RNA-seq data of the snMultiome analysis were normalized and scaled using SCTransform v2 (v0.4.1) in Seurat (v5.1.0). Cell cycle effects (G2M vs. S phase) and mitochondria genes were scored and regressed out prior to integration. Quality-filtered gene-by-nucleus matrices were then integrated across samples using reciprocal PCA in Seurat, following established best practices.

For snMultiome ATAC data, a consensus peak set was generated across all six samples, resulting in 252,290 ATAC peaks used as variable features for fragment counting and integration. Peak-by-nucleus count matrices were integrated using reciprocal latent semantic indexing (LSI) projections via the Signac package (v1.14.0).

### Dimensional reduction, clustering and cell type annotation

We applied a weighted nearest neighbor (WNN) analysis in Seurat, integrating 1–30 RNA principal components and 2–50 ATAC latent semantic indexing (LSI) components to capture multimodal structure. The resulting WNN graph was used for UMAP visualization and clustering. Cluster identities were assigned based on canonical marker gene expression. To further resolve progenitor diversity, clusters 0, 2, 5, 9, 11, 16, 26, 27, and 29 were extracted for secondary dimensional reduction and clustering. Only genes identified as variable by SCTransform and detected in more than 10% of cells within a cluster were tested as marker genes. MAST was used to identify cluster-enriched genes (|logFC| > 0.25, p < 0.01). Cell-type annotations were curated based on established markers from previous studies (Figure 1).

### Marker peaks and peak annotation

We identified marker peaks—ATAC-seq peaks showing cell type–specific accessibility—using the FindAllMarkers function in Seurat v5 and performed logistic regression (LR) tests within the union peak set. To account for technical variability, total fragment number was included as a latent variable in the model, mitigating the effects of differential sequencing depth across cells, as suggested by Ntranos et al. (2018) for single-cell RNA-seq data. deepTools (v3.5.6) was used to visualize the top 1,000 marker peaks per cluster centered on peak summits using the plotHeatmap function. Functional enrichment analysis of cluster-specific peak sets was performed using the GREAT (v4.04) online tool. The whole genome was used as background, and the “basal plus extension” rule was applied to associate genomic regions with nearby genes. In GREAT, each gene is assigned a basal regulatory domain of 5 kb upstream and 1 kb downstream of its transcription start site (TSS), which is extended in both directions to the nearest gene’s basal domain, up to 1 Mb in either direction. Functional terms meeting the criteria of binomial FDR < 0.05, hypergeometric FDR < 0.05, and region-based fold enrichment ≥ 2 were retained. Among these, biologically relevant terms were selected and visualized based on their binomial FDR ranking.

### Gene regulatory network analysis

To infer gene regulatory networks during human GE development, we applied the SCENIC+ (1.0a2) framework to our single-nucleus multiome data, with all inputs prepared according to SCENIC+ guidelines [PMID: 37443338]. For this dataset, topic modeling was performed with pycisTopic [PMID: 30962623] using default parameters, and 100 topics were selected as optimal based on log-likelihood. A custom cistarget database was constructed from a consensus peak set of 252,290 peaks, using the SCENIC+ human v10 motif collections. eRegulon matrices were subsequently used for downstream analysis.

### Trajectory analysis

To reconstruct developmental trajectories of the basal ganglia, we extracted cells corresponding to the LGE and MGE GABAergic lineages from the full dataset and analyzed them with Slingshot (v2.10.0) [PMID: 29914354]. Each lineage consisted of its RG, IPCs, neuroblasts, and associated neuronal subtypes—LGE immature inhibitory neurons and MSNs for the LGE, and MGE interneurons, MGE_SST, and MGE_CRABP1 for the MGE. For trajectory reconstruction, we ensured region specificity by keeping only the progenitors from the same region (LGE RG/IPCs/neuroblasts and MGE RG/IPCs/neuroblasts). For both LGE and MGE lineages, the weighted nearest-neighbour graph (1–30 SCT PCs; 2–30 LSI) was recalculated and used to generate the UMAP. To delineate the overall lineage architecture, we built a minimum spanning tree at the cluster level. We anchored the tree by assigning the GE-RG cluster as the starting node and marked clusters representing fully differentiated cell types as terminal endpoints. To capture the continuous developmental progression, we fitted principal curves to each inferred lineage. Each cell was assigned a weight reflecting its projection distance to the curve. Pseudotime was then derived from the curve ordering, and shrinkage was applied to individual branches to ensure smooth convergence of the trajectories.

To characterize temporal changes in eRegulon activity, we used tradeSeq (v1.16.0) to fit generalized additive models that map eRegulon AUC scores onto pseudotime for each trajectory [PMID: 32139671]. We employed beta-regression generalized additive models with six knots in tradeSeq, since AUC scores are bounded between 0 and 1. Fitted trajectories were obtained with predictSmooth across 100 evenly spaced pseudotime positions per lineage. Ten eRegulon modules per trajectory were identified by k-means clustering on the fitted AUC values for LGE and MGE.

For functional annotation, we performed GO Biological Process enrichment analysis using the one-sided hypergeometric test in clusterProfiler (v4.10.1). For each module, genes detected in ≥8% of constituent eRegulons were defined as core target genes and served as the query gene set. The background consisted of the union of all eRegulon target genes. Enrichment results for all modules are provided in DataSX

### Comparative Analysis of Cortical and GE Radial Glia

MetaNeighbor analysis [PMID: 34234317] was used to quantify similarities between cortical and GE progenitor subtypes using highly variable genes or peaks. A pretrained model was generated from second-trimester cortical progenitor reference data [PMID: 39131371]. GE progenitor profiles were then evaluated against this reference using the MetaNeighborUS algorithm. Cell-type similarity was measured using area under the receiver operating characteristic curve (AUROC) scores, with higher values indicating greater transcriptomic or epigenomic similarity across datasets. To visualize similarity and difference between GE RG subtypes and their most similar cortical RG subtypes, AUROC values were z-score transformed within each cell type for visualization.

As an independent approach, we quantified similarity between cortical and GE RG subtypes by computing Spearman correlations using the average expression of marker genes or accessibility of marker peaks that were differentially expressed or differentially accessible in either dataset (log₂FC ≥ 0.25, adjusted P ≤ 0.01).

To detect differentially expressed genes and differentially accessible peaks between cortical and GE RG, we generated pseudobulk profiles by aggregating expression or accessibility values by sample and region. DESeq2 [PMID: 25516281] was used to evaluate differential expression and accessibility on genes or peaks with ≥1% expression or accessibility.

### CellChat analysis

Cell-cell communication was inferred using CellChat (v2.2.0) [[PMID: 39289562]] with default settings. Batch-corrected expression profiles were supplied to CellChat in two separate analyses: one using all 20,319 nuclei and another using only the progenitor subset. Communication probabilities were computed across all curated ligand–receptor pairs in CellChatDB, retaining only those for which at least ten cells in the sending or receiving populations expressed the ligand or receptor. The resulting communication network was consolidated at the signaling pathway level and represented as a weighted, directed graph, with weights corresponding to the aggregated communication probabilities between cell types. Network structure was visualized using chord diagrams, where outer bars indicate signaling sources, inner bars indicate signaling targets with sizes proportional to the magnitude of incoming signal, and edge thickness reflects interaction strength.

### SCAVENGE

We used SCAVENGE (v1.0.2)40 to map disease-associated genetic variation onto single-nucleus chromatin accessibility profiles [PMID: 35668323]. We considered six neuropsychiatric disorders (ASD, MDD, BPD, AHDH, OCD, and SCZ). The GWAS summary statistics used in this study were downloaded from the following sources:

ADHD: https://figshare.com/articles/dataset/adhd2022/22564390 ASD: https://figshare.com/articles/dataset/asd2019/14671989 MDD: https://datashare.ed.ac.uk/handle/10283/3203

SCZ: http://figshare.com/articles/dataset/scz2022/19426775 BPD: https://figshare.com/articles/dataset/bip2024/27216117 OCD: https://figshare.com/articles/dataset/ocd2024/28707155

ABF finemapping (prior = 1×10⁻⁴) was performed on 1 Mbp windows around GCTA-COJO–derived sentinel variants (P < 5×10⁻⁸) [PubMed ID: 22426310, PubMed ID: 21167468]. Summary statistics were filtered for MAF > 0.01. Variants were ranked by posterior probability to define 95% credible sets.

Finemapped posterior probabilities from ABF finemapping were imported into gchromVAR to calculate trait-relevant scores (TRS) for each cell. A cell-by-peak count matrix was constructed from integrated single-nucleus ATAC-seq data with GC bias correction. Network propagation and TRS scaling and normalization were performed using default parameters [PMID: 35668323]. Cells in the top 0.1% of TRS scores were considered trait-relevant, with statistical significance determined through 1,000 permutations (P < 0.001). Cell type enrichment was evaluated using two-sided hypergeometric tests with Benjamini-Hochberg correction; cell types with FDR < 0.01 and odds ratio > 1.4 were considered significantly enriched for trait-associated variants.

### Spatial transcriptomics analysis

An automated cell segmentation on our MERFISH datasets was performed as described in a published study^12^. A custom neural network model was trained on manually segmented images incorporating both nuclear and cell boundary stainings. Vizgen post-processing analytical tools were used to generate the cell-by-gene expression matrix, along with spatial and image intensity metadata at the single-cell level. The following quality control criteria was applied to the single-cell data for experiments on two technical replicates from a GW23 individual (batch 1) and one section from a GW14 individual (batch 2): a minimum transcript count of 25 (15 for batch 2), and a maximum of 2,000; detection of at least 10 genes; and an estimated cell volume between 100 and three times the median across all cells. Data normalization, variance stabilization, and scaling were carried out using SCTransform (v0.4.1^13^). Clustering was performed using Louvain algorithm and cell type annotation was determined based on the expression of canonical markers and differentially expressed genes. Identification of spatial homogeneous regions was achieved through a joint analysis of spatial coordinates and cell type labels, employing a k-nearest neighbor graph-based approach (Jackson et al., 2024). To evaluate how a given cell type influences the expression of neighboring cells in the MGE region containing DENs structures, we performed an intercellular communication modelling analysis consisting of a graph neural network-based approach to model gene expression profiles as a function of cell type annotations and cell neighborhoods^14^. Sender-receiver effects were evaluated according to the magnitude of the fold change in log-scale as well as the significance of the FDR-corrected p-values (Benjamini-Hochberg correction) obtained through a Wald test on the parameter estimates of the sender-receiver interaction matrix.

## Quantifications and Statistical analysis

### Live Imaging Analysis

Acquired time-lapse images were maximum intensity projected and merged if separate files had been acquired per position (due to resetting of z position/focus). Division types and MST translocation distances were analysed using the manual spots function in Imaris. To be considered MST, a “jump” of the cell body along a single prominent fibre was required. MST length was manually tracked within the spots function. At least three separate imaging positions across at least 2 individual slices were analysed per biological sample. Quantifications were conducted using Imaris Software.

### Quantification

Quantifications of cell numbers (immunostaining) were conducted using Imaris Software. For statistical analysis, an unpaired two-tailed student’s t-test using prism software was used.

For DEN quantifications in PCDH19 KO lines and isogenic controls, DCX-regions that were Nestin+ or vimentin+ (and not a rosette/lumen) were quantified in a blinded manner.

### Division angle analysis

Time-lapse images were opened with Fiji Image J software. The plane of division was measured using the three-point built-in angle measurement tool of the software in relationship to primary fibre orientation. At least two individual slices with at least 5 cells each were processed per biological sample.

### Statistics

The average of all technical replicates per biological sample was calculated and used for statistical testing. Outliers were removed using a ROUT test (Q = 5%) in prism. For statistical analysis, an unpaired, parametric, two-tailed student’s t-test assuming both populations have the same standard deviation (SD) using prism software was used.

